# Regulation of parvalbumin interneuron plasticity by neuropeptide-encoding genes

**DOI:** 10.1101/2023.02.03.527010

**Authors:** Martijn Selten, Clémence Bernard, Fursham Hamid, Alicia Hanusz-Godoy, Fazal Oozeer, Christoph Zimmer, Oscar Marín

## Abstract

Neuronal activity is regulated in a narrow permissive band for the proper operation of neural networks. Changes in synaptic connectivity and network activity, for example, during learning, might disturb this balance, eliciting compensatory mechanisms to maintain network function. In the neocortex, excitatory pyramidal cells and inhibitory interneurons exhibit robust forms of stabilising plasticity. However, while neuronal plasticity has been thoroughly studied in pyramidal cells, little is known about how interneurons adapt to persistent changes in their activity. Here we uncover the critical cellular and molecular mechanisms through which cortical parvalbumin-expressing (PV+) interneurons adapt to changes in their activity levels. We found that changes in the activity of PV+ interneurons drive cell-autonomous, bi-directional compensatory adjustments of the number and strength of inhibitory synapses received by these cells, specifically from other PV+ interneurons. High-throughput profiling of ribosome-associated mRNA revealed that increasing the activity of PV+ interneurons leads to the cell-autonomous upregulation of two genes encoding multiple secreted neuropeptides, *Vgf* and *Scg2*. Functional experiments demonstrated that VGF is critically required for the activity-dependent scaling of inhibitory PV+ synapses onto PV+ interneurons. Our findings reveal an instructive role for neuropeptide-encoding genes in regulating synaptic connections among PV+ interneurons in the adult mouse neocortex.

---

Stabilising network activity is essential for the operation of neural circuits, most critically during experience-dependent plasticity and learning (Miller and MacKay, 1994; Turrigiano, 2011; Barth and Ray, 2019; Humeau and Choquet, 2019). Neurons regulate their activity through several mechanisms, including altering their intrinsic excitability and modulating the strength and number of synapses they receive (Turrigiano et al., 1994, 1998; Kirkwood et al., 1996; Davis and Goodman, 1998; Desai et al., 1999; Burrone et al., 2002; Maffei et al., 2006). The precise cellular and molecular mechanisms through which neurons undergo synaptic plasticity are, to a large extent, cell type-specific (Turrigiano, 2012; Hooks and Chen, 2020).

In the cerebral cortex, the activity of individual neurons depends on the balance between synaptic excitation and inhibition, which is mediated by glutamatergic pyramidal cells and gamma-aminobutyric acid-containing (GABAergic) interneurons, respectively. Pyramidal cells receive stable ratios of excitation and inhibition because any increase in excitation leads to a proportional increase in inhibition through the recruitment of inhibitory inputs (Vogels et al., 2011; Xue et al., 2014). This activity-dependent reorganisation of inhibitory connections targeting pyramidal cells requires changes in gene expression and is mediated by retrograde signals from the pyramidal cells (Peng et al., 2010; Bloodgood et al., 2013; Yap et al., 2021). Although interneurons likely exhibit similar forms of plasticity (Spiegel, 2020), the extreme diversity of cortical GABAergic cells has so far prevented their characterisation. In particular, how individual interneurons cell-autonomously adapt to changes in their activity remains largely unknown.

Fast-spiking basket cells expressing the calcium-binding protein parvalbumin (PV+) are the most abundant type of GABAergic interneuron in the mouse neocortex (Markram et al., 2004). PV+ interneurons play a pivotal role in maintaining the excitation-inhibition balance of cortical circuits through potent perisomatic inhibition of pyramidal cells (Yazaki-Sugiyama et al., 2009; House et al., 2011; Kuhlman et al., 2013; Hu et al., 2014). PV+ interneurons regulate experience-dependent plasticity (Reh et al., 2020) and are involved in forming neuronal assemblies during learning (Letzkus et al., 2011; Donato et al., 2013). While it is evident that PV+ interneurons exhibit robust plasticity (Chang et al., 2010; Ruediger et al., 2011; Karunakaran et al., 2016; Favuzzi et al., 2017), the cellular mechanisms and molecular signals regulating the adaptation of cortical PV+ interneurons to activity changes remain poorly characterised.

## Homeostatic scaling of inhibitory connectivity is bidirectional

Most research on stabilising plasticity in the neocortex has relied on sensory deprivation paradigms, which makes it challenging to disentangle cell-autonomous from non-cell-autonomous mechanisms due to the complexity of the network compensations that follow these experimental manipulations. To overcome this limitation, we used a chemogenetic approach based on Designer Receptor Exclusively Activated by Designer Drugs (DREADDs), which allows transient neuronal activation (hM3Dq) or inhibition (hM4Di) following administration of the pharmacologically inert molecule clozapine-N-oxide (CNO) (Roth, 2016), targeting specifically neocortical PV+ interneurons. In brief, we injected layer 2/3 of the primary somatosensory cortex (S1) of neonatal *Pvalb^Cre/+^* mice with an adeno-associated virus (AAV) encoding Cre-dependent hM3Dq and mCherry to restrict its expression to PV+ interneurons (Fig. 1a). Most critically, we titrated the virus to infect only a relatively small fraction of PV+ interneurons in each animal, thereby minimising the possibility of indirect, network-induced changes (Supplementary Fig. 1). This approach also allowed us to monitor non-cell-autonomous changes in neighbouring PV+ interneurons not infected by the virus. We then treated young adult mice with CNO or vehicle for two days and examined the synaptic inputs received by hM3Dq-expressing (hM3Dq+) PV+ interneurons using whole-cell electrophysiology. As a proxy of their activation, we measured Fos protein levels and found that hM3Dq+ PV+ cells contained higher levels of Fos in mice treated with CNO than in control mice (Fig. 1b,c). We also found that the amplitude and frequency of miniature excitatory postsynaptic currents (mEPSCs) recorded from hM3Dq+ PV+ interneurons were not affected by the transient activation of these cells (Fig. 1d-e). In contrast, both the amplitude and frequency of miniature inhibitory postsynaptic currents (mIPSCs) were increased following the activation of PV+ interneurons (Fig. 1f-g), which led to an overall reduction in the excitatory/inhibitory (E/I) ratio of these cells (Fig. 1h). This effect was caused by CNO-mediated activation of hM3Dq+ PV+ interneurons, as CNO treatment in mice injected with mCherry-expressing (mCherry+) AAVs did not affect the amplitude or frequency of mEPSCs and mIPSCs (Supplementary Fig. 2).

**Fig. 1.**
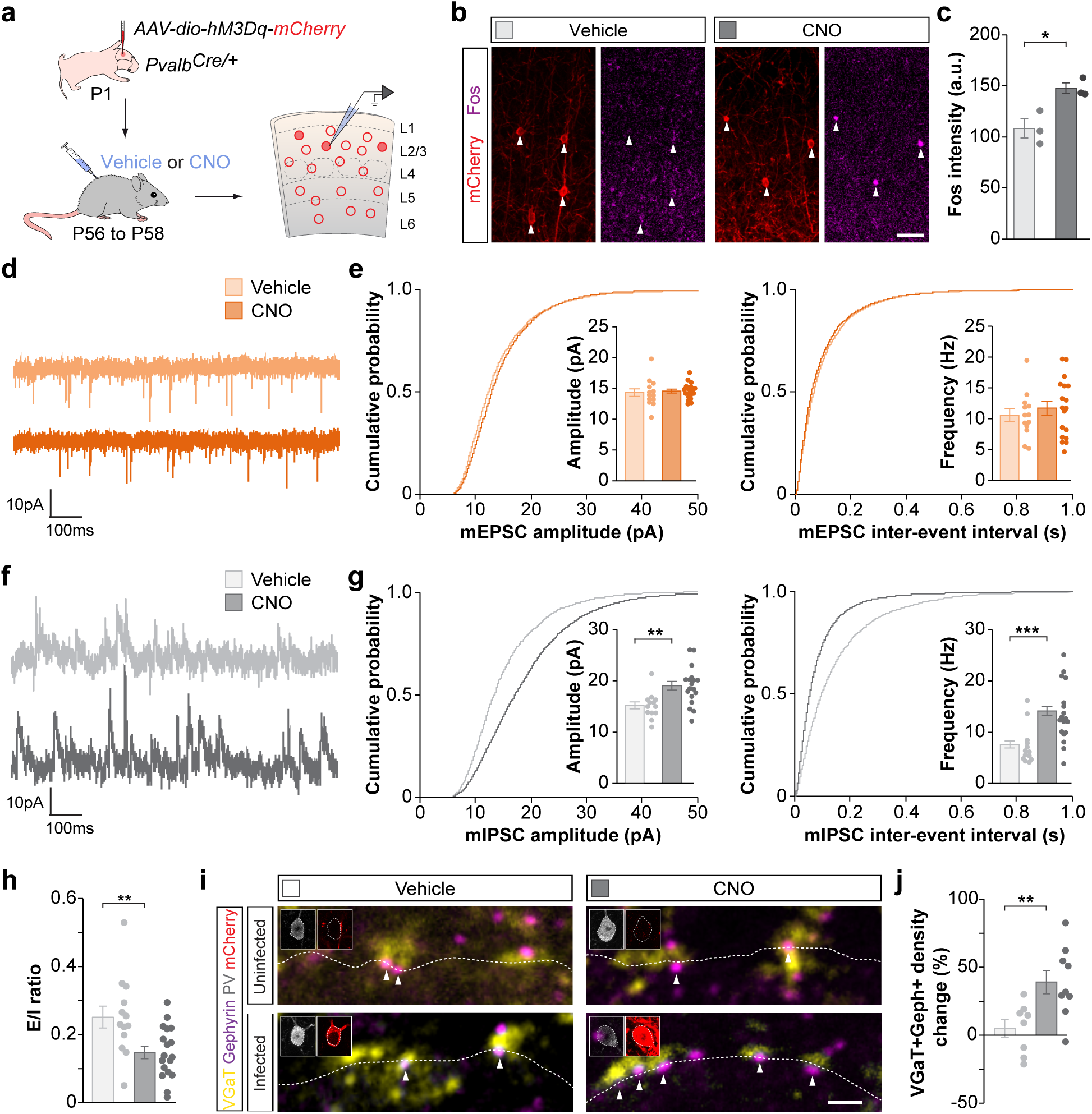
| Inhibitory synapses mediate the homeostatic response of PV+ interneurons to increased activity. **a**, Experimental strategy. **b**, Fos and PV expression in coronal sections through S1. **c**, Quantification of the intensity of Fos staining in hM3Dq-mCherry infected PV+ interneurons in vehicle- and CNO-treated mice (vehicle, *n* = 3 mice; CNO, *n* = 3 mice; two-tailed Student’s t-test, *p* = 0.02). **d**, Traces of mEPSCs recorded from hM3Dq-mCherry infected PV+ interneurons in vehicle- and CNO-treated mice. **e**, Quantification of amplitude (vehicle, *n* = 13 cells; CNO, *n* = 18 cells; two-tailed Student’s t-test, *p* = 0.71) and frequency (vehicle, *n* = 13 cells; CNO, *n* = 18 cells; two-tailed Student’s t-test, *p* = 0.48) of mEPSCs recorded from hM3Dq-mCherry infected PV+ interneurons in vehicle- and CNO-treated mice. **f**, Traces of mIPSCs recorded from hM3Dq-mCherry infected PV+ interneurons in vehicle- and CNO-treated mice. **g**, Quantification of amplitude (vehicle, *n* = 13 cells; CNO, *n* = 18 cells; two-tailed Student’s t-test, *p* = 0.003) and frequency (vehicle, *n* = 13 cells; CNO, *n* = 18 cells; Mann-Whitney U-test, *p* = 0.001) of mIPSCs recorded from hM3Dq-mCherry infected PV+ interneurons in vehicle- and CNO-treated mice. **h**, Excitation/Inhibition (E/I) ratio in hM3Dq-mCherry infected PV interneurons in vehicle- and CNO-treated mice (vehicle, *n* = 13 cells; CNO, *n* = 18 cells; two-tailed Student’s t-test, *p* = 0.005). **i**, Presynaptic VGaT+ puncta and postsynaptic Gephyrin+ clusters in infected and uninfected PV+ interneurons in vehicle- and CNO-treated mice. Inserts show PV and mCherry immunoreactivity in cell somas. **j,** Quantification of the change in synaptic density between infected and uninfected PV+ interneurons in vehicle- and CNO-treated mice (vehicle, *n* = 8 mice; CNO, *n* = 9 mice; two-tailed Student’s t-test, *p* = 0.008). Data are presented as mean ± s.e.m. Scale bar, 50 μm (b) and 1 μm (k).

To investigate whether an increase in inhibitory synapses accompanied the elevated inhibitory drive observed in activated PV+ interneurons, we performed immunohistochemistry against pre-(VGaT) and postsynaptic (Gephyrin) markers of inhibitory synapses and quantified their density on the soma of hM3Dq+ PV+ interneurons. We observed an increase in the density of inhibitory synaptic clusters contacting hM3Dq+ PV+ interneurons compared to uninfected neighbouring PV+ interneurons in CNO-treated but not vehicle-treated mice (Fig. 1i,j and Supplementary Fig 3). These experiments revealed that the increase in inhibitory drive observed in hM3Dq+ PV+ interneurons is accompanied by an increase in the number of inhibitory synapses received by these cells. Moreover, this effect is cell-autonomous, as only activated PV+ interneurons showed increased synapse density.

We next investigated whether the inhibitory inputs received by PV+ interneurons change bidirectionally in response to changes in the activity of these cells. To this end, we reduced the activity of PV+ interneurons by expressing a Cre-dependent inhibitory DREADD (hM4Di; Supplementary Fig. 4a). Following CNO administration, we observed no changes in the amplitude and frequency of mEPSCs recorded from hM4Di-expressing PV+ interneurons (Supplementary Fig. 4c,d). In contrast, reducing the activity of PV+ interneurons led to a decrease in the frequency of mIPSCs recorded from these cells (Supplementary Fig. 4e,f), which resulted in an overall increase in the E/I ratio (Supplementary Fig. 4b). Altogether, these experiments revealed that modulating the activity of PV+ interneurons in vivo leads to a bidirectional and cell-autonomous compensatory change in the inhibitory drive these cells receive.

Changes in the number of inhibitory inputs dynamically follow the activity of PV+ cells. Do these inputs originate from a specific interneuron population? PV+ interneurons receive synapses from the three largest subclasses of interneurons: other PV+ interneurons, somatostatin-expressing (SST+) interneurons, and vasoactive intestinal peptide-expressing (VIP+) interneurons (Pfeffer et al., 2013; Jiang et al., 2015). To determine the contribution of each of these populations to the changes in presynaptic inhibitory input that follow the increased activity of PV+ interneurons, we used optogenetic stimulation to evoke synaptic output from each subclass of interneuron (PV+, SST+ or VIP+) while recording from hM3Dq+ PV+ interneurons (Fig. 2a). To this end, we first generated mice expressing Flippase (Flp) recombinase under the *Pvalb* locus along with Cre-dependent channelrhodopsin (ChR2) alleles (*Pvalb^Flp/Flp^*;*RCL^ChR2/Chr2^*), and then crossed these mice with *Pvalb^Cre/+^*, *Sst^Cre/+^* and *Vip^Cre/+^* mice to generate animals in which ChR2 was expressed in one of each of the three subclasses of interneurons (*Pvalb^Cre/Flp^;RCL^ChR2/+^*, *Sst^Cre/+^;Pvalb^Flp/+^;RCL^ChR2/+^*, and *Vip^Cre/+^;Pvalb^Flp/+^;RCL^ChR2/+^*). To modify the activity of PV+ interneurons in these experiments, we injected these mice with AAVs encoding a Flp-dependent hM3Dq (Fig. 2a). As in previous experiments, we titrated the virus to infect a small fraction of PV+ interneurons (Supplementary Fig. 5a,b) and verified its effectiveness to activate hM3Dq+ PV+ interneurons using Fos (Supplementary Fig. 5c,d).

**Fig. 2.**
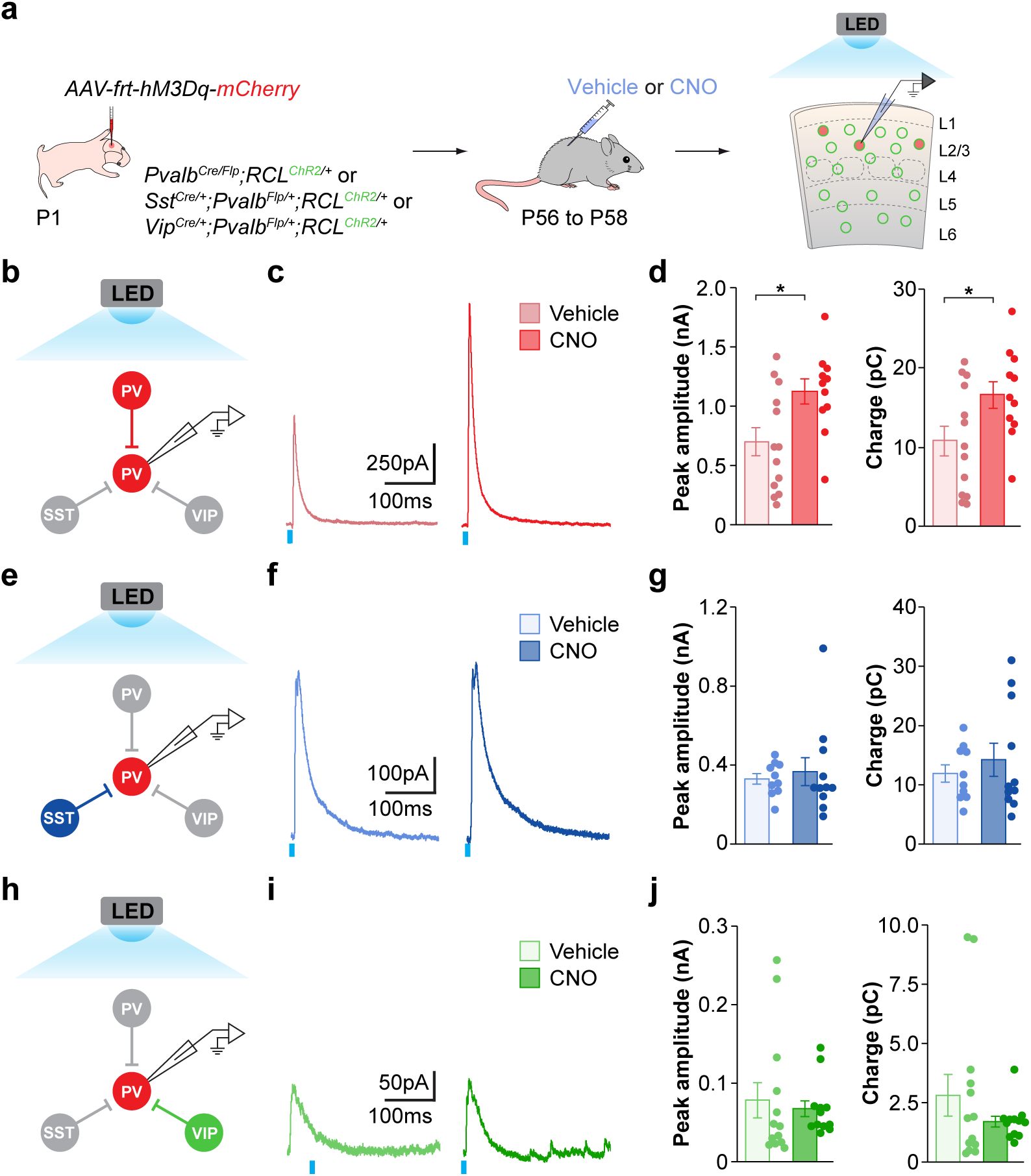
| The homeostatic inhibition originates from PV+ interneurons. **a**, Experimental strategy. **b**, Recording and stimulation configuration in *Pvalb^Cre/Flp^;RCL^Chr2/+^* mice. **c**, Traces of eIPSCs recorded from hM3Dq+ PV+ interneurons following full-field stimulation of PV+ interneurons in vehicle- and CNO-treated mice. **d,** Peak amplitude (vehicle, *n* = 13 cells; CNO, *n* = 11 cells; two-tailed Student’s t-test, *p* = 0.02) and charge (vehicle, *n* = 13 cells; CNO, *n* = 11 cells; two-tailed Student’s t-test, *p* = 0.04) of LED-evoked eIPSCs. **e**, Recording and stimulation configuration in *Sst^Cre/+^;Pvalb^Flp/+^;RCL^Chr2/+^* mice. **f**, Traces of eIPSCs recorded from hM3Dq+ PV+ interneurons following full-field stimulation of SST+ interneurons in vehicle- and CNO-treated mice. **g**, Peak amplitude (vehicle, *n* = 10 cells; CNO, *n* = 11 cells; two-tailed Student’s t-test, *p* = 0.64) and charge (vehicle, *n* = 10 cells; CNO, *n* = 11 cells; two-tailed Student’s t-test, *p* = 0.48) LED-evoked eIPSCs. **h**, Recording and stimulation configuration in *Vip^Cre/+^;Pvalb^Flp/+^;RCL^Chr2/+^* mice. **i**, Traces of eIPSCs recorded from hM3Dq+ PV+ interneurons following full-field stimulation of VIP+ interneurons in vehicle- and CNO-treated mice. **j**, Peak amplitude (vehicle, *n* = 13 cells; CNO, *n* = 12 cells; two-tailed Student’s t-test, *p* = 0.67) and charge (vehicle, *n* = 13 cells; CNO, *n* = 12 cells; two-tailed Student’s t-test, *p* = 0.25) LED-evoked eIPSCs. Data are presented as mean ± s.e.m.

We first examined the contribution of other PV+ interneurons to the increased inhibitory drive observed in hM3Dq+ PV+ interneurons following their increased activity (Fig. 2b). Because ChR2 is also expressed in the recorded cells from *Pvalb^Cre/Flp^;RCL^ChR2/+^* mice, we performed control whole-cell patch-clamp recordings from synaptically isolated PV+ interneurons to confirm that the contribution of ChR2-mediated currents is negligible compared to the evoked synaptic currents in our recording conditions (Supplementary Fig. 6). We tested if CNO treatment affected the excitability of both infected and uninfected PV+ interneurons in response to ChR2 stimulation (Supplementary Fig. 7a-f). While infected PV+ interneurons in CNO-treated mice exhibited reduced excitability at higher LED stimulus intensities, stimulation at 10% LED power (7.5 mW/mm^2^) led to the same number of evoked action potentials as in infected PV+ interneurons from vehicle-treated mice (Supplementary Fig. 7e,f). We recorded spontaneous inhibitory postsynaptic currents (sIPSCs) from each cell before the ChR2 stimulation to verify the increase in inhibitory inputs following treatment with CNO. As expected from previous experiments, we observed increased amplitude and frequency of sIPSC in hM3Dq+ PV+ interneurons from CNO-treated compared to vehicle-treated *Pvalb^Cre/Flp^;RCL^ChR2/+^* mice (Supplementary Fig. 8a,b). ChR2-mediated stimulation of PV+ interneurons also revealed an increase in the peak amplitude and charge of ChR2-evoked IPSCs recorded from hM3Dq+ PV+ interneurons in CNO-treated mice compared to controls (Fig. 2c,d). Importantly, this increase was not due to global presynaptic strengthening of all PV+ synapses because we detected no changes in sIPSCs or ChR2-evoked synaptic currents when recording from pyramidal cells (Supplementary Fig. 8c,d and Supplementary Fig. 9). These experiments demonstrated that the increased inhibition received by PV+ interneurons following their prolonged activation derives, at least in part, from other PV+ interneurons. To corroborate this observation, we quantified the number of synapses made by PV+ interneurons on hM3Dq+ PV+ interneurons and uninfected neighbouring cells using the PV+ specific presynaptic marker synaptotagmin-2 (Syt2) (Sommeijer and Levelt, 2012). We found an increase in the density of PV+ synaptic clusters contacting hM3Dq+ PV+ interneurons compared to uninfected neighbouring PV+ interneurons in CNO-treated but not vehicle-treated mice (Supplementary Fig. 10). Altogether, these experiments revealed that increasing the activity of PV+ interneurons leads to a cell-autonomous, homeostatic increase in the number of inhibitory synapses these cells receive from other PV+ interneurons.

We next tested whether SST+ and VIP+ interneurons also contribute to the increased inhibitory drive observed in hM3Dq+ PV+ interneurons following increased activity using the same strategy (Fig. 2a). First, we verified that CNO treatment did not affect the excitability of SOM+ or VIP+ interneurons (Supplementary Fig. 7g-l). However, we observed no changes between CNO-treated mice and controls when recording ChR2-evoked synaptic currents in hM3Dq+ PV+ interneurons following optogenetic stimulation of SST+ or VIP+ interneurons (Fig. 2e-j). sIPSCs recorded from the same cells revealed an increase in amplitude and frequency in CNO-treated mice from both *Sst^Cre/+^;Pvalb^Flp/+^;RCL^ChR2/+^* and *Vip^Cre/+^;Pvalb^Flp/+^;RCL^ChR2/+^* mice, which confirmed that in these experiments CNO treatment also led to an increase in inhibitory inputs contacting hM3Dq+ PV+ interneurons (Supplementary Fig. 8e-h). Together, these experiments demonstrated that raising the activity of PV+ interneurons increases the strength and number of inhibitory inputs these cells receive from other PV+ interneurons but not from SST+ or VIP+ interneurons.

## Activity-dependent gene expression in PV+ interneurons

To identify the molecular mechanisms through which PV+ interneurons modulate the inputs they receive from other PV+ interneurons in response to changes in their activity, we obtained the translatome of hM3Dq+ PV+ interneurons in control and CNO-treated mice. To this end, we used a viral approach to access ribosome-associated mRNAs by translating ribosome affinity purification (Nectow et al., 2017). In brief, we engineered Flp-dependent AAVs allowing the bicistronic expression of hemagglutinin (HA)-tagged ribosomal protein (Rpl10a) and myc-tagged hM3Dq. Because PV is transiently expressed in some layer 5 pyramidal cells, we also flanked this construct with LoxP sites so that Cre-mediated recombination would prevent its expression. We injected this virus into S1 of *Pvalb^Flp/+^*;*Neurod6^Cre/+^* mice to achieve specific and sparse expression in PV+ interneurons (Supplementary Fig. 11a,b) and verified its effectiveness to activate hM3Dq+ PV+ interneurons using Fos (Supplementary Fig. 11c,d). We then isolated S1 from mice treated with vehicle or CNO for 48 h, pulled-down ribosomes using anti-HA beads to isolate ribosome-associated mRNA transcripts from PV+ interneurons, and quantified expression levels by RNA sequencing (Fig. 3a).

**Fig. 3.**
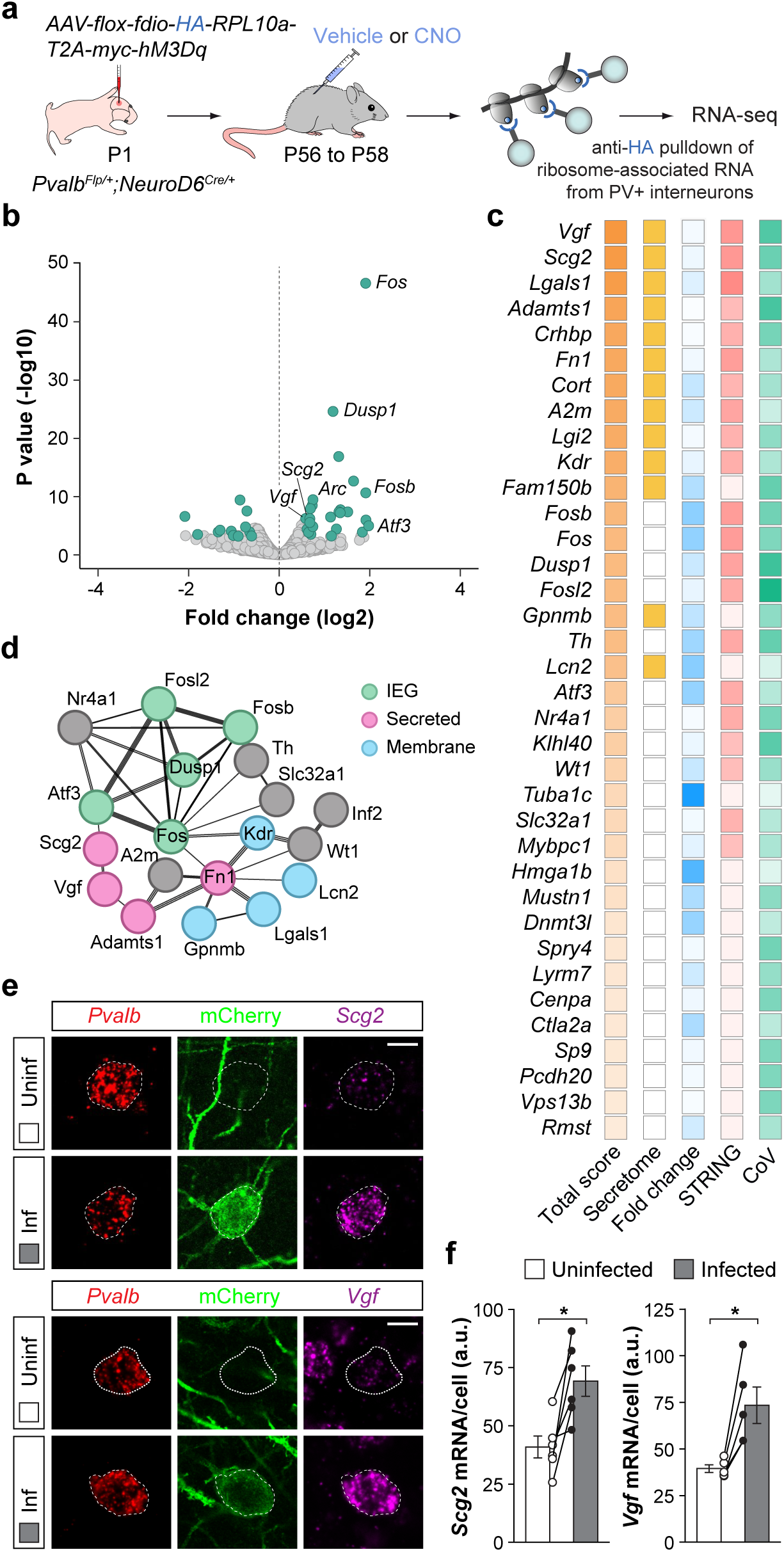
| Increased activity in PV+ interneurons leads to the upregulation of *Scg2* and *Vgf*. **a**, Experimental strategy. **b,** Volcano plot showing ribosome-associated mRNAs identified through RNA-sequencing. Differentially expressed RNAs (FC > 1.5, P < 0.05) are labelled in teal. **c,** Ranking of the top DEGs based on four selection criteria (see Methods). Darker colours indicate higher scores (values 0 to 1). **d,** Partial STRING network of the most prominent node of DEGs, highlighting IEGs, secreted and membrane-bound proteins. **e,** *Scg2* and *Vgf* expression in neighbouring hM3Dq+ and uninfected PV+ interneurons in layer 2/3 of S1 from CNO-treated mice. **f,** Quantification of the expression of *Scg2* (*n* = 6 mice; two-tailed paired Student’s t-test, *p* = 0.01) and *Vgf* (*n* = 5 mice; two-tailed paired Student’s t-test, *p* = 0.01) in neighbouring hM3Dq+ and uninfected PV+ interneurons from CNO-treated mice. Data are presented as mean ± s.e.m. CoV, coefficient of variation. Scale bar, 10 μm.

We found 51 differentially expressed genes (DEGs), 37 upregulated and 14 downregulated, in PV+ interneurons from vehicle- and CNO-treated mice (Fig. 3b). Among the upregulated genes, we identified several immediate early genes (IEGs) such as *Fos*, *FosB*, and *Fosl2*, which is consistent with the increased activity of hM3Dq+ PV+ interneurons treated with CNO. We ranked DEGs according to several criteria, including fold expression change, biological reproducibility, the strength of predicted protein-protein interactions, and the secretory nature of the proteins (Fig. 3c, see Methods). The top candidates on this prioritised list included *Adamts1*, which encodes a disintegrin and metalloproteinase with thrombospondin motif protein family member responsible for cleaving lecticans in the perineuronal nets surrounding PV+ interneurons (Gottschall and Howell, 2015), *Lgals1*, which encodes a glycoprotein of the galectin family developmentally enriched in PV+ basket cells (Favuzzi et al., 2019), and two neuropeptide-encoding genes, *Scg2* (Secretogranin II) and *Vgf* (non-acronymic, also known as *Scg7*), which have been linked to synaptic plasticity (Alder et al., 2003; Yap et al., 2021). Protein-protein interaction network analysis by STRING (Szklarczyk et al., 2020) revealed that many DEGs constitute an active node enriched in IEGs and other transcription factors, along with the two top candidates, *Scg2* and *Vgf* (Fig. 3d). We confirmed that *Scg2* and *Vgf* mRNA expression was upregulated in a cell-autonomous manner following the activation of PV+ interneurons in vivo using single-molecule fluorescence in situ hybridisation (Fig. 3e, f and Supplementary Fig. 12). Together, these results revealed that cortical PV+ interneurons react to a sustained increase in their activity by inducing the expression of a molecular programme that includes IEGs, genes linked to the remodelling of perineuronal nets, and genes encoding secretable neuropeptides.

## Vgf mediates inhibitory plasticity in PV+ interneurons

To test whether *Scg2* and *Vgf* mediate the changes in PV-PV synaptic connectivity during plasticity, we sought to downregulate the expression of these genes, specifically in activated PV+ interneurons. To this end, we designed Cre-dependent AAVs expressing both hM3Dq and a short hairpin RNA (shRNA) against the candidate genes (*shScg2, shVgf*, and *shLacZ* as a control) and validated their ability to downregulate the expression of the target genes (Supplementary Fig. 13a-c). We then performed three separate experiments injecting *Pvalb^Cre/+^* mice with AAVs conditionally expressing hM3Dq and shRNAs against *Scg2* (*shScg2-hM3Dq*), *Vgf* (*shVgf-hM3Dq*), or *LacZ* (*shLacZ-hM3Dq*). As in previous experiments, viral injections were titrated to infect only a small population of PV+ interneurons and confirmed that the three vectors led to the activation of infected PV+ interneurons following CNO administration (Supplementary Fig. 13d-f).

We assessed the effect of downregulating *Scg2* and *Vgf* expression in the formation of PV+ synapses contacting other PV+ interneurons following treatment with vehicle or CNO (Fig. 4a). We found no changes in the density of PV+ (Syt2+/Gephyrin+) synapses contacting the soma of neighbouring infected and uninfected PV+ interneurons in vehicle-treated mice. This result suggested that the prolonged downregulation of *Scg2* and *Vgf* alone does not modify the basal inhibitory connectivity of PV+ interneurons (Supplementary Fig. 14). We then examined whether the downregulation of *Scg2* or *Vgf* would interfere with the plasticity of inhibitory synapses contacting PV+ interneurons in CNO-treated mice. As expected, we observed a substantial increase in the density of Syt2+/Gephyrin+ puncta contacting infected PV+ interneurons in mice injected with *shLacZ-hM3Dq* and treated with CNO (Fig. 4b,c).

**Fig. 4.**
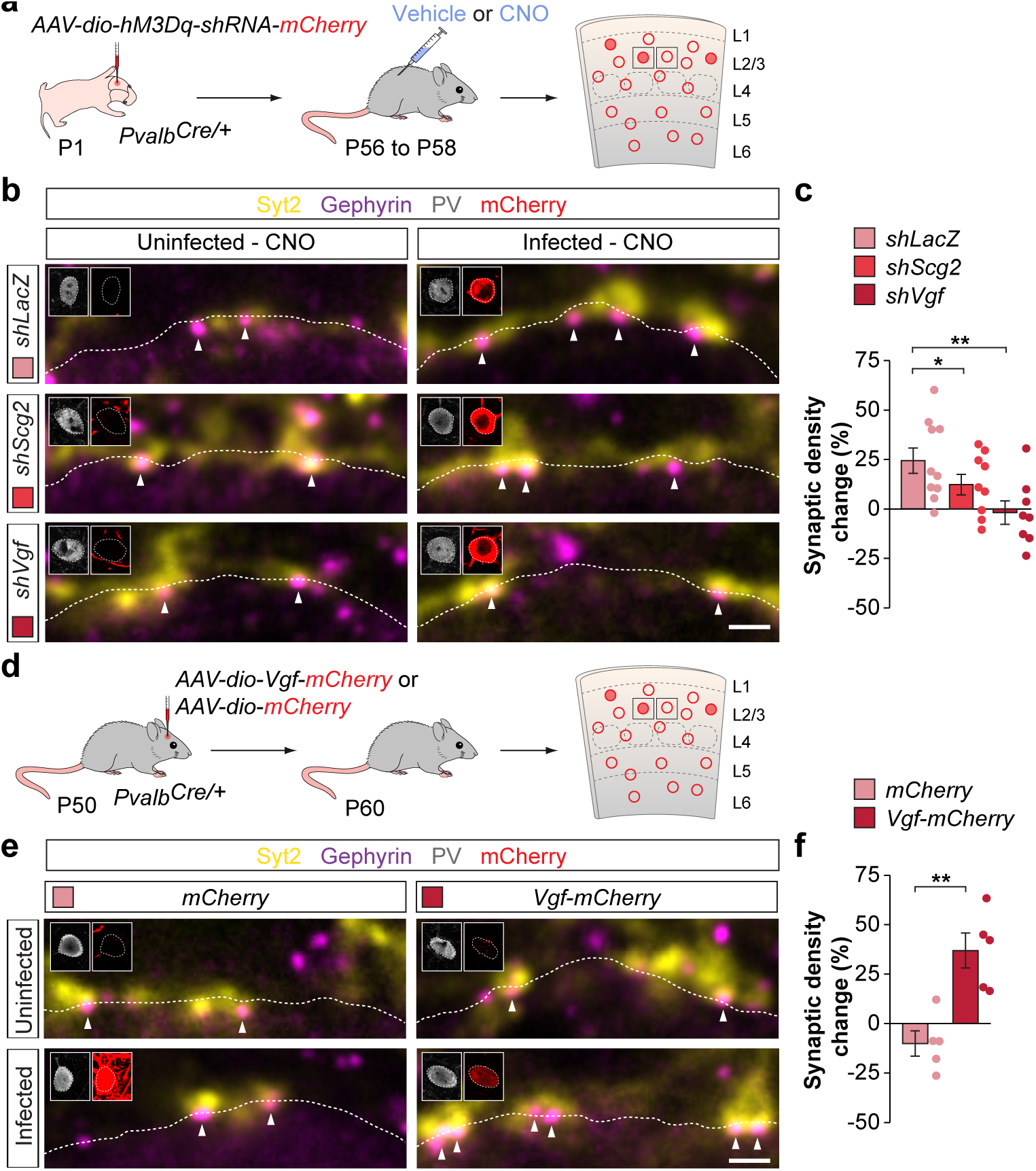
| VGF is necessary and sufficient for scaling PV-PV connectivity. **a**, Experimental strategy. **b**, Presynaptic Syt2+ puncta and postsynaptic Gephyrin+ clusters in *shLacZ, shScg2* and *shVgf* expressing PV+ interneurons in vehicle- and CNO-treated mice. Inserts show PV and mCherry immunoreactivity in cell somas. **c**, Quantification of the change in synaptic density between infected and uninfected PV+ interneurons in CNO-treated mice injected with *hM3Dq-shLacZ* (*n* = 10 mice), *hM3Dq-shScg2* (*n* = 9 mice, one-sample t-test, *p* = 0.049) and *hM3Dq-shVgf* (*n* = 8 mice, one-sample t-test, *p* = 0.003). **d,** Experimental strategy. **e**, Presynaptic Syt2+ puncta and postsynaptic Gephyrin+ clusters in *mCherry-* and *Vgf-mCherry* expressing PV+ interneurons. Inserts show PV and mCherry immunoreactivity in cell somas. **f,** quantification of change in synaptic density between neighbouring uninfected and infected PV+ interneurons in mice injected with *Vgf-mCherry* (uninfected, *n* = 5 mice; CNO, *n* = 5 mice; two-tailed Student’s t-test, *p* = 0.003). Data are presented as mean ± s.e.m. Scale bars, 1 μm.

This effect was attenuated in mice injected with *shScg2-hM3Dq* and completely abolished in mice injected with *shVgf-hM3Dq* (Fig. 4b,c and Supplementary Fig. 14c-h). In other words, while we continued to observe an increase in PV+ synapse density on PV+ interneurons infected with *shLacZ* following CNO treatment, PV+ interneurons infected with *shScg2* had a reduced ability to increase the number of PV+ synapses they received in response to an increase in their activity, whereas PV+ cells infected with *shVgf* were completely unable to increase the number of PV+ synapses they receive. These experiments demonstrated that the upregulation of *Scg2* and, most notably, *Vgf* in PV+ interneurons following their activation is required for the formation of new afferent synapses from other PV+ interneurons in response to changes in their activity.

We then asked if increased expression of *Vgf* is sufficient for the increase in PV-PV connectivity in the absence of increased activity. To this end, we generated Cre-dependent AAVs to overexpress full-length VGF and mCherry or mCherry alone, injected them into S1 of P50 *Pvalb^Cre/+^*mice, and assessed changes in inhibitory synaptic density 10 days after injection (Fig. 4d and Supplementary Fig. 15a-c). We found a significant increase in the density of PV+ synapses targeting PV+ interneurons infected with Vgf-mCherry compared to uninfected neighbouring control cells (Fig 4e,f and Supplementary Fig. 15f,g). In contrast, we did not observe this increase in PV+ interneurons infected only with mCherry (Fig. 4e,f and Supplementary Fig. 15d-e). These experiments demonstrated that increased expression of Vgf, in the absence of increased activity, is sufficient to increase PV-PV interconnectivity.

## Discussion

Our results shed light on a fundamental mechanism mediating the efficient balance of excitation and inhibition in the cerebral cortex. PV+ interneurons balance the excitation received by pyramidal cells by dynamically adjusting their inhibitory inputs to these cells (Xue et al., 2014). We demonstrate here that PV+ interneurons are also responsible for regulating the activity of other PV+ interneurons through the modulation of their mutual inhibitory synapses. Connections among cortical PV+ interneurons are more frequent than among any other type of interneuron or between other interneurons and PV+ interneurons (Somogyi et al., 1998; Galarreta and Hestrin, 1999; Gibson et al., 1999; Avermann et al., 2012; Bezaire and Soltesz, 2013). This reciprocal connectivity may allow PV+ interneurons to control their firing rate and synchronise with each other during gamma oscillations (Bartos et al., 2007). In addition, PV+ interneurons are frequently self-connected by autapses – synapses that a neuron makes onto itself– which allow them to adjust their temporal firing interval with extreme accuracy (Deleuze et al., 2019; Szegedi et al., 2020). The dynamic regulation of the interconnectivity of PV+ interneurons likely influences their modulation of gamma-rhythms (Bartos et al., 2007; Buzsáki and Wang, 2012), which are essential for sensory perception and attention.

The activity-dependent adjustment of inhibition in PV+ interneurons involves changes in gene expression, a process likely coordinated by members of the Fos family of transcription factors (Cohen et al., 2016). In the hippocampus, Fos-dependent changes in the expression of *Scg2* in pyramidal cells modulate the inhibition these cells receive from PV+ interneurons (Yap et al., 2021). Although Scg2 also seems to be involved in the regulation of inhibitory synapses among PV+ interneurons, our study reveals that VGF is the critical factor regulating the plasticity of these synapses. These observations suggest that neuropeptide-encoding genes may have evolved to modulate the activity-dependent plasticity of different inhibitory connections in the cerebral cortex.

VGF is a neuropeptide precursor of the extended granin family. These proteins are stored and sorted into dense core vesicles, proteolytically processed, and secreted as small bioactive peptides (Bartolomucci et al., 2011). Several secreted neuropeptides derived from VGF, including TLQP-62 and TLQP-21, have been previously linked to memory formation in the hippocampus and energy homeostasis in the hypothalamus (Bartolomucci et al., 2006; Lin et al., 2015), although their mechanism of action remains unclear. Determining the specific VGF-derived peptides involved in synapse formation, their pre- and postsynaptic sites of action, and the putative G-protein coupled receptor mediating their function in PV+ interneurons will be critical for establishing the precise mechanisms through which VGF regulates inhibitory synaptic plasticity in these cells.

Dysregulation of VGF expression has been associated with multiple neurodegenerative and psychiatric conditions, most notably in Alzheimer’s disease and mood disorders (Quinn et al., 2021). Alterations in PV+ interneurons may mediate functional deficits in many of these conditions (Marín, 2012; Contractor et al., 2021; Exposito-Alonso and Rico, 2022). Identifying a critical role for VGF in regulating synaptic connectivity among PV+ interneurons provides a path to mechanistically understand its contribution to the plasticity of cortical circuits in health and disease.

## Acknowledgements

We thank E. Serafeimidou-Pouliou for laboratory support, M. Moissidis for help with script writing and I. Andrew for managing mouse colonies, Silvia Arber (*Pvalb^Cre/+^*) and Klaus Nave (*Nex^Cre/+^*) for mice, and Bryan Roth, Bong-Kiun Kaang, and Liqun Luo for plasmids. We are grateful to J. Burrone, M. Grubb, T. Kroon, and B. Rico for critical reading of the manuscript and to members of the Marín and Rico laboratories for stimulating discussions and ideas. This work was supported by grants from the European Research Council (ERC-2017-AdG 787355) to O.M. and the Rosetrees Trust (PGL21/10159) to O.M. and M.S.

## Author contributions

M.S. and O.M. designed the experiments. M.S. performed most experiments, except for the biochemical purification of mRNAs carried out by C.B. F.H. performed the bioinformatic analyses. A.H-G. and C.Z. contributed to data collection and analysis. F.O. produced the AAVs. M.S. and O.M. wrote the manuscript.

## Competing interests

The authors declare no competing interests.

## Materials and Methods

### Mice

#### Mouse lines and breeding

All experiments were performed following King’s College London Biological Service Unit’s guidelines and the European Community Council Directive of November 24, 1986 (86/609/EEC). Animal work was carried out under licence from the UK Home Office following the Animals (Scientific Procedures) Act 1986. Similar numbers of male and female mice were used in all experiments. Animals were maintained under standard laboratory conditions on a 12:12 light/dark cycle with water and food ad libitum. The *Pvalb^Cre/+^* mice used in most experiments were generated by crossing *Pvalb^Cre/+^* mice (*B6.129P2-Pvalb^tm1(cre)Arbr^/J*, JAX017320) with CD1 mice (Crl:CD1[ICR], Charles River). To generate *Pvalb^Cre/Flp^;RCL^ChR2/+^*, *Sst^Cre/+^;Pvalb^Flp/+^;RCL^ChR2/+^* and *Vip^Cre/+^;Pvalb^Flp/+^;RCL^ChR2/+^* mice, we first generated *Pvalb^Flp/Flp^;RCL^ChR2/ChR2^* mice by crossing *Pvalb^Flp/Flp^* mice (*B6.Cg-Pvalb^tm4.1(flpo)Hze^/J*, JAX022730) with *RCL^ChR2/ChR2^* (Ai32) mice (*B6;129S-Gt(ROSA)26Sor^tm32(CAG-COP4*H134R/EYFP)Hze/^J*, JAX012569). These mice were then crossed with *Pvalb^Cre/Cre^*, *Sst^Cre/Cre^* (*Sst^tm2.1(cre)Zjh^/J*, JAX013044) or *Vip^Cre/Cre^* (*Vip^tm1(cre)Zjh^/J*, JAX010908) mice. *Pvalb^Flp/+^;NeuroD6^Cre/+^* mice were generated by crossing *Pvalb^Flp/+^* mice with *Nex^Cre/+^* (*Neurod6^tm1(cre)Kan^*) mice (Goebbels et al., 2006).

#### CNO treatment

Mice between P42 and P70 were treated with vehicle or Clozapine N-oxide (CNO, Tocris Cat. No. 4936) as indicated. Mice used for Fos experiments received a single injection of vehicle or CNO (1 mg/kg) and were perfused 2 h later. In all other experiments, mice were treated with vehicle or CNO (1 mg/kg) for ∼48h via 5 injections. Injections were administered starting 2 days before the experimental day and took place in the morning (between 8 am and 10 am) and in the evening (between 6 pm and 9 pm). A fifth injection was administered between 8 am and 10 am on the day of the experiment. CNO was dissolved in 0.5% Dimethyl Sulfoxide (DMSO, Sigma, D0418) in 0.9% saline and stored at -20 °C.

### Histology

#### Immunohistochemistry

Mice were deeply anaesthetised with pentobarbital before being transcardially perfused with 0.9% saline, followed by 4% paraformaldehyde (PFA) in PBS. Brains were postfixed for 2h in 4% PFA at 4°C, followed by cryoprotection in 15% and 30% sucrose. Brains were sectioned at 40 µm on a sliding microtome (Leica SM2010R) and stored in ethylene glycol (30% ethylene glycol (Merck, 324558), 30% Glycerol (Merck, G5516) in PBS) at -20 °C. Free-floating sections were washed in 0.25% Triton-X100 (Merck) in phosphate-buffered saline (PBS) before 2 h incubation in blocking buffer containing 0.25% Triton-X100, 10% Serum and 2% Bovine Serum Albumin in PBS. Sections were incubated overnight at 4 °C in a blocking buffer with primary antibodies. The next day, sections were washed in 0.25% Triton-X100 in PBS and incubated for 2h in a blocking buffer with secondary antibodies at room temperature (RT). Sections were then washed in PBS, and stained with 5 µM 4′,6-diamidino-2-phenylindole (DAPI) (Merck) in PBS if required. Sections were dried and mounted with Mowiol/DABCO (8% Mowiol (SLS, 81381), 2% Dabco (Merck, D27802) in PBS) . The following primary antibodies and concentrations were used: Goat anti-mCherry (1:500, Antibodies-Online, ABIN1440057), dsRed anti-rabbit (1:500, Clontech, 632496), c-Fos anti-rabbit (1:200, Merck, ABE457), Gephyrin anti-Mouse-IgG1 (1:500, Synaptic Systems, 147011), VGaT anti-Guinea pig (1:500, Synaptic Systems, 131004), Parvalbumin anti-Chicken (1:250, Synaptic Systems, 195006), Parvalbumin anti-Mouse (1:3000, Swant, 235), Synaptotagmin-2 anti-Mouse-IgG2 (1:250, ZFIN, ZDB-ATB-081002-25). The following secondary antibodies and concentrations were used: Donkey anti-Chicken 405 (1:200, Jackson, 703-475-155), Goat anti-Mouse-IgG1 488 (1:500, Molecular Probes, A21121), Donkey anti-Rabbit 488 (1:400, Thermo Fisher Scientific, A21206), Donkey anti-Rabbit 555 (1:400, Molecular Probes, A31572), Donkey anti-Goat 555 (1:400, Invitrogen, A21432), Goat anti-Mouse-IgG2 647 (1:500, Molecular Probes, A21241), Donkey anti-Guinea Pig 647 (1:250, Jackson, 706-605-148), Donkey anti-Mouse IgG1 647 (1:400, Thermo Fisher Scientific, A31571).

#### Single-molecule fluorescence in situ hybridization

Brains were perfused, sectioned and stored in RNase-free solutions. Mice were perfused as described above and postfixed for 24h before being cryoprotected in 15% and 30% sucrose. The brains were sectioned at 30 µm on a sliding microtome and stored at 20 °C. Sections were mounted on RNase-free SuperFrost Plus slides (ThermoFisher) and probed following the RNAscope Multiplex Fluorescent Assay v2 protocol (ACDBio #323110). The following probes from ACDBio were used: Mm-Scg2-C1 (#477691), Mm-Vgf-C1 (#517421) and Mm-Pvalb-C2 (#421931).

#### Image acquisition

Images were acquired using an inverted SP8 confocal microscope using the LAS AF software. Different experimental conditions within an experiment were always imaged in parallel, and imaging settings were kept constant for each experiment. All images were taken at 200 Hz acquisition speed and 1024×1024-pixel resolution. Infection density and Fos intensity were imaged using a 10x objective (0.8800 pixels/μm). RNAscope images were acquired using a 63x objective (5.5440 pixels/μm). Synaptic quantification images were taken using a 100x objective at 1.75x digital zoom (15.4000 pixels/μm). Imaging was restricted to S1 L2/3 unless otherwise stated. To image neighbouring cells, we aimed to image an infected and an uninfected cell as close as possible together at the same distance from the pial surface (e.g., after imaging an infected PV+ interneuron, an effort was made to image an uninfected PV+ interneuron nearby and at a similar depth from the pial surface).

#### Image analysis

Fos intensity was analysed using a custom MATLAB script. Cell bodies were segmented using disk morphological shape function, size, and intensity thresholding to create individual regions of interest (ROIs). L2/3 was determined using DAPI counterstaining. ROIs in L2/3 were used to measure average signal intensity in the Fos channel.

For single-molecule fluorescence in situ hybridization experiments, the number of mRNA particles was determined using a custom MATLAB script. Briefly, for each image, background subtraction was applied. The contour of PV+ cells was then used to draw an ROI and labelled as mCherry+ or mCherry-after visual inspection. Within each ROI, the total area of the signal was measured. This area is then divided by the area of a single mRNA particle (∼0.16 µm^2^), which estimates the number of mRNA particles in the ROI. We applied intensity correction to normalise signal intensity differences between brains. For this, we measured the average signal intensity of each image, excluding the ROI of the PV+ interneurons and calculated the average signal intensity for each brain. We normalized these values to the lowest value to get the intensity ratio. We then divided the number of mRNA particles by the intensity ratio.

Synaptic densities were analysed using a custom FIJI script, as previously described earlier (Favuzzi et al., 2017). Background subtraction, Gaussian blurring, smoothing, and contrast enhancement were applied in all channels. Cell somas were drawn automatically or manually based on intensity levels of Parvalbumin staining to create a mask of the soma surface and measure its perimeter. Presynaptic boutons and postsynaptic clusters were detected automatically based on thresholds of intensity. Thresholds for the different synaptic markers were selected from a set of random images before quantification, and the same threshold was applied to all images from the same experiment. The “Analyze Particles” and “Adjustable Watershed” tools were applied to the synaptic channels, and a mask was generated with a minimum particle size of 0.05. The soma mask and the corresponding synaptic masks were merged to quantify the number of puncta contacting the soma and dendrite. Puncta were defined as presynaptic boutons when they were located outside the soma or dendrite and had ≥ 0.1 µm^2^ colocalizing with the soma or dendrite perimeter. Puncta were defined as postsynaptic clusters contained inside a soma or dendrite and had ≥ 0.2 µm^2^ colocalizing with the soma or dendrite perimeter. Synapses were defined as presynaptic boutons and postsynaptic clusters contacting each other with a colocalization area of ≥ 0.03 µm^2^ of their corresponding masks. Classification of cells as uninfected or infected was performed manually based on the absence or presence of mCherry signal, respectively. The density change was calculated as:

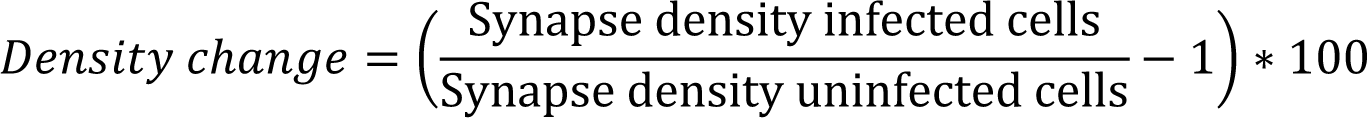

### Viruses

#### Virus production

AAV8 viruses were produced in HEK293FT cells grown on 5 plates (linear PEI, Polysciences Europe Cat. No. 23966-100) or 10 plates (branched PEI, Sigma-Aldrich Cat. No. 408727) of 15 cm diameter until they reached 60% confluency. Cells were grown on DMEM (Gibco 21969-035) + 10% fetal bovine serum (FBS) (Gibco 10500-064) + 1% penicillin/streptomycin (Gibco 15140-122) and 10mM HEPES. AAVs were produced using Polyethylenimine (PEI) transfection of HEK293FT cells with a virus-specific transfer plasmid (70µg/10 plates) and a pDP8.ape helper plasmid (300µg/10 plates, PF478 from PlasmidFactory). The helper plasmid provided the AAV Rep and Cap functions and the Ad5 genes (VA RNAs, E2A and E4). The DNA and PEI were mixed in a 1:4 ratio in uncomplemented DMEM and left at room temp for 25 mins to form the DNA:PEI complex. The transfection solution was added to each plate and incubated for 72 h at 37°C in 5% CO_2_. The transfected cells were then scraped off the plates and pelleted. The cell pellet was lysed in lysis buffer containing 50 mM Tris-Cl, 150 mM NaCl and 2 mM MgCl_2_ and 0.5% Sodium deoxycholate and incubated with 100 U/ml Benzonase (Sigma, E1014 25KU) for 1hr to dissociate particles from membranes. The particles were cleared by centrifugation, and the clear supernatant was filtered through 0.8 µm (Merck Millipore Ref SLAA033SS) and 0.45 µm (Merck Millipore Ref SLHA 033SS) filters. The viral suspension was loaded on a discontinuous iodixanol gradient using four layers of different iodixanol concentrations (Zolotukhin et al., 1999) of 15, 25, 40 and 58% in Quick-seal polyallomer tubes (Beckman Coulter, Cat. No. 342414) and spun in a VTi-50 rotor at 50.000 rpm for 1h 15 min at 12 °C in an Optima L-100 XP Beckman Coulter ultracentrifuge to remove any remaining contaminants. After completion of the centrifugation, 5 ml were withdrawn from the 40/58% interface using a G20 needle. The recovered virus fraction was purified by first passing through a 100-kDA molecular weight cut-off (MWCO) centrifugal filter (Sartorius VIVASPIN VS2041) and then through an Amicon Ultra 2ml Centrifugal filter (Millipore UFC210024). Storage buffer (350 mM NaCl and 5% Sorbitol in PBS) was added to the purified virus, and 5 µl aliquots were stored at -80 °C.

The following viruses were used in this study: *AAV8-hSyn-DIO-hM3D(Gq)-mCherry* (Addgene 44361), *AAV8-hSyn-DIO-mCherry* (Addgene 50459), *AAV8-hSyn-DIO-hM4D(Gi)-mCherry* (Addgene 44362), *AAV8-hSyn-fDIO-hM3D(Gq)-mCherry*, *AAV8-hSyn-flx-fDIO-HA-Rpl10a-T2A-myc-hM3D(Gq), AAV8-hSyn-DIO-hM3D(Gq)-mCherry-shLacZ-CWB* (*shRNA* sequence: AAATCGCTGATTTGTGTAGTC), *AAV8-hSyn-DIO-hM3D(Gq)-mCherry-shScg2-CWB* (*shRNA* sequence: GCAGACAAGCACCTTATGAA), *AAV8-hSyn-DIO-hM3D(Gq)-mCherry-shVgf-CWB* (*shRNA* sequence: GACGATCGATAGTCTCATTGA), *AAV8-hSyn-DIO-Vgf-mCherry-CW3SL*. *AAV8-hSyn-DIO-hM3D(Gq)-mCherry*, *AAV8-hSyn-DIO-mCherry* and *AAV8-hSyn-DIO-hM4D(Gi)-mCherry* were a gift from Bryan Roth (Krashes et al., 2011). *pAAV-CWB-EGFP* and *pAAV-CW3SL-EGFP* backbones were used to create plasmids as indicated and were a gift from Bong-Kiun Kaang (Choi et al., 2014). *pAAV-hSyn-FLExFRT-mGFP-2A-Synaptophysin-mRuby* was used to create fDIO plasmids and was a gift from Liqun Luo (Beier et al., 2015). To achieve sparse infection, all viruses were diluted to a titre between 5×10^11^ and 8×10^11^.

### Stereotactic injections

#### Juvenile injection

P1-3 pups were anaesthetized with isoflurane for juvenile injections and mounted on a stereotactic frame (Stoelting). Mice used for vTRAP/RNAseq experiments received six 150 nl injections at a speed of 10 nl/sec in S1 in the left hemisphere. All other mice received three 150 nl injections at a speed of 10 nl/sec in S1 in the left hemisphere. For vTRAP experiments, viruses were supplemented with green fluorescent beads (LumaFluor) to label the injection sites. Mice were allowed to recover in a heated recovery chamber (Vettech, HEO11) before returning to the home cage.

#### Adult injections

P42-60 mice were anaesthetized with isoflurane and mounted on a stereotactic frame. A small incision was made in the skin over the injection area on the right hemisphere, and the skull was cleaned using a cotton swab. Two holes were drilled using a rotary drill (Foredom, K.1070) at the following coordinates (from Bregma): (1) anteroposterior +1.0 mm and mediolateral -3.2 mm; and (2) anteroposterior +1.5 mm and mediolateral -3.2 mm. Mice received two 300 nl injections at a speed of 100 nl/min at each location. After injection, the injection capillary was left in place for three minutes before retraction. Following injections, the skin was sutured with absorbable vicryl sutures (Ethicon, W9500T), and mice were allowed to recover in a heated recovery chamber at 36 °C before being returned to the home cage.

### Electrophysiology

#### Slice preparation

Mice (P42-70) were deeply anesthetized with an overdose of sodium pentobarbital and transcardially perfused with 10 mL ice-cold N-Methyl-D-glucamine (NMDG) solution containing (in mM) 93 NMDG, 2.5 KCl, 1.2 NaH_2_PO_4_, 30 NaHCO_3_, 20 HEPES, 25 glucose, 5 sodium ascorbate, 2 thiourea, 3 sodium pyruvate, 10 MgSO_4_ and 0.5 CaCl_2_ (300-310 mOsmol, pH 7.3-7.4) oxygenated with 95% O_2_ and 5% CO_2_. Following decapitation, the brain was quickly removed, and the injected hemisphere was glued to a cutting platform before submerging in ice-cold NMDG solution. Coronal slices (300 µm) were cut using a vibratome (VT1200S, Leica) and placed in NMDG solution at 32 °C for 11 minutes before being transferred to a holding solution containing (in mM) 92 NaCl, 2.5 KCl, 1.2 NaH_2_PO_4_, 30 NaHCO_3_, 20 HEPES, 25 glucose, 5 sodium ascorbate, 2 thiourea, 3 sodium pyruvate, 2 MgSO_4_ and 2 CaCl_2_ (300-310 mOsmol, pH 7.3-7.4), oxygenated with 95% O_2_ and 5% CO_2_, at room temperature for at least 45 minutes until recording. All salts were purchased from Sigma-Aldrich.

#### Electrophysiological recordings

Slices were transferred to the recording setup 15 min prior to the recording while being continuously superfused with recording ACSF containing (in mM) 124 NaCl, 1.25 NaH_2_PO_4_, 3 KCl, 26 NaHCO_3_, 10 Glucose, 2 CaCl_2_, and 1 MgCl_2_, continuously oxygenated with 95% O_2_ and 5% CO_2_ at 32 °C. Pipettes (3–5 MΩ) were made from borosilicate glass capillaries using a PC-10 pipette puller (P10, Narishige) and filled with intracellular solution containing (in mM) 115 CsMeSO_3_, 20 CsCl, 10 HEPES, 2.5 MgCl_2_, 4 Na_2_ATP, 0.4 Na_3_GTP, 10 sodium-phosphocreatine, 0.6 EGTA (pH 7.2–7.3, 285–295 mOsm). Cells were visualised with an upright microscope (Olympus) and recorded using a Multiclamp 700B amplifier (Molecular Devices). The signal was passed through a Hum Bug Noise Eliminator (Digitimer), sampled at 20 kHz, and filtered at 3 kHz using a Digidata 1440A (Molecular devices). All cells were recorded in L2/3 of S1bf. Miniature excitatory postsynaptic currents (mEPSCs) and miniature inhibitory postsynaptic currents (mIPSCs) were recorded in the presence of 1 μM Tetrodotoxin (Tocris, 1069) at a holding voltage of -60 mV and +10mV, respectively. In order to record mEPSCs and mIPSCs from the same cells, synaptic currents were not blocked. Cells were excluded if the access resistance (Ra) exceeded 25 MΩ or I_hold_ > 200 pA (at V_hold_ = -60mV). For ChR2 evoked IPSCs (eIPSCs), slices were incubated in recording ACSF supplemented with 5 μM 6-Cyano-7-nitroquinoxaline-2,3-dione (CNQX, Tocris, 1045) and 100 μM D-(−)-2-Amino-5-phosphonopentanoic acid (D-APV, Tocris, 0106) to block glutamatergic inputs. Cells were held at a holding voltage of + 10mV, and after allowing the cell to settle (approximately 1 min), sIPSCs were recorded. Subsequently, whole-cell series resistance was 70% compensated. eIPSC were evoked with full field 5 ms LED pulses (power: PV-PV 10% LED, SOM-PV 10% LED, VIP-PV 100% LED, as indicated for the respective experiments. 100% LED power = 75.1 mW/mm^2^) through a 4x objective (Olympus) with an inter-stimulus interval of 60 s from a pE100 illumination system (CoolLED). Cells were excluded from analysis if the series resistance changed > 20%, Ra > 25 MΩ or I_hold_ > 200 pA (at V_hold_ = -60mV), and only cells with both sIPSC and eIPSC recordings were included in the data. Cell-attached recordings were performed using recording electrodes filled with recording ACSF in the presence of CNQX and D-APV, loose seals (around 200 MΩ) were formed on cells, and LED stimulus of various intensities was applied via a 4x objective in the same way as in the eIPSC recordings (see above).

#### Electrophysiology analysis

Miniature (mEPSC and mIPSC) and spontaneous (sIPSC) recordings were analysed using Mini Analysis (Synaptosoft). E/I ratio was calculated (excitatory current/second)/(inhibitory current/second). Current/second was calculated as (average charge of an event) x frequency. E/I ratios were calculated for each recorded cell, and only cells with mEPSC and mIPSC recordings that passed quality control (see above) were included. eIPSC and cell-attached recordings were analysed in Clampfit 10.2 (Molecular Devices).

### Biochemistry and RNA sequencing

#### RNA isolation by anti-HA pulldown

The infected cortex from P44 – P70 mice, injected with *AAV8-hSyn-flx-fDIO-HA-Rpl10a-T2A-myc-hM3D(Gq)* and treated for 48 h with vehicle or CNO, was rapidly dissected in ice-cold RNase-free PBS, using fluorescent beads as a guide to identifying the infected area, and immediately homogenised in ice-cold homogenisation buffer (50 mM Tris-HCl pH 7.5, 100 mM KCl, 12 mM MgCl2, 1 mg/ml Heparin [Sigma-Aldrich], cOmplete EDTA-free protease inhibitors [Sigma-Aldrich], 200 U/ml RNAsin [Promega], 100 µg/ml cycloheximide and 1 mM DTT [Sigma-Aldrich]). Tissue from 4 brains was pooled for every sample. Samples were centrifuged at 2,000 g for 10 min at 4°C, and Igepal-CA630 (Sigma-Aldrich) was added to the samples to a final concentration of 1%. Samples were then centrifuged at 13,000 g for 10 min at 4°C and the supernatant was added on 100 µl of anti-HA magnetic beads (Pierce #88837, previously washed in homogenisation buffer) for 3-4 h at 4°C with gentle rotation. After incubation, beads were washed 3 times in ice-cold washing buffer (300 mM KCl, 1% Igepal-CA630, 50 mM Tris-HCl pH 7.5, 12mM MgCl2, 1mM DTT and 100 µg/ml cycloheximide) and eluted in 350 µl of RLT Plus buffer from the RNAeasy Plus Micro kit (Qiagen) supplemented with 2-mercaptoethanol (Bio-Rad).

#### RNA purification, quantification, and quality check

RNA purification of immunoprecipitated RNA was performed using the RNeasy Plus Micro kit (Qiagen) following the manufacturer’s protocol. RNA quality was checked on a Bioanalyzer instrument (Agilent Technologies) using an RNA 6000 Pico Chip. Only RNA samples with RNA integrity number (RIN) values higher than 9 were used for library preparation and sequencing.

#### Library preparation and Illumina sequencing

Four biological replicates were analysed for each genotype. The Genomic Unit of the Centre for Genomic Regulation (CRG, Barcelona, Spain) performed the library preparation and RNA sequencing. The library was prepared using the SMARTer Ultra Low RNAkit and samples were then sequenced paired-end using an Illumina HiSeq 2500 platform to a mean of approximately 60 million mapped reads per sample.

#### Bioinformatics

High-throughput sequencing data from PV+ interneurons-derived ribosome-associated mRNAs were processed using the community-curated Nextflow (version 21.03.0.edge, build 5518 [05-03-2021 10:52 UTC], available at https://zenodo.org/record/3490660#.Y8AhHXbP2Uk) RNA-seq pipeline(Ewels et al., 2020). Specifically, sequencing reads were quality-controlled by FastQC (available at https://www.bioinformatics.babraham.ac.uk/projects/fastqc/) and quality-trimmed by Trim Galore (available at https://zenodo.org/record/5127899#.Y8fdOi-l3UI). Mouse GRCm38/mm10 genome annotation was accessed from Illumina’s iGenomes repository (available at https://support.illumina.com/sequencing/sequencing_software/igenome.html) and used as a reference for read alignment by STAR(Dobin et al., 2013) and for gene abundance quantification by Salmon(Patro et al., 2017). Gene-level counts data were imported into R using the tximport package(Soneson et al., 2016) and analysed by edgeR(Robinson et al., 2010) using the estimateGLMRobustDisp model. Genes with less than 10 reads in at least 4 samples were excluded from the analysis(Chen et al., 2016). The > 1.5-fold, FDR < 0.05 and TpM > 1 cut-off were used to identify genes that respond to PV+ interneuron activation (Supplementary Table S1). Activity-dependent genes were ranked using four scoring criteria: fold-change, coefficient of variation (CoV), STRING, and secretome scores. The fold-change score indicates the magnitude of the response and is scaled for a value between 0 and 1. The CoV score represents the degree of reproducibility and was calculated by scaling the sum of the ratio of the standard deviation to the mean from each sample. The STRING score of each gene signifies its co-interaction with other activation-response genes and was computed by scaling the sum of interaction scores obtained from STRING enrichment analysis (Szklarczyk et al., 2020). The secretome score takes up a binary value. Genes that are part of the human secretome (Uhlén et al., 2019) were assigned a value of 1.

### Statistical analyses

All statistical analyses were performed using SPSS (Supplementary Table S2). No statistical methods were used to predetermine sample sizes. Sample sizes were chosen based on previous publications in the field. Experimental mice from all genotypes or conditions were processed together. Samples were tested for normality using the Shapiro–Wilk normality test. Differences were considered significant when p values < 0.05. Data are presented as mean ± s.e.m. Statistical details of experiments are described in figure legends.

### Data availability

RNA-seq data will be deposited into the public repository Gene Expression Omnibus (GSE223038) upon publication. All other data will be shared upon request.

---

**Supplementary Fig. 1.**
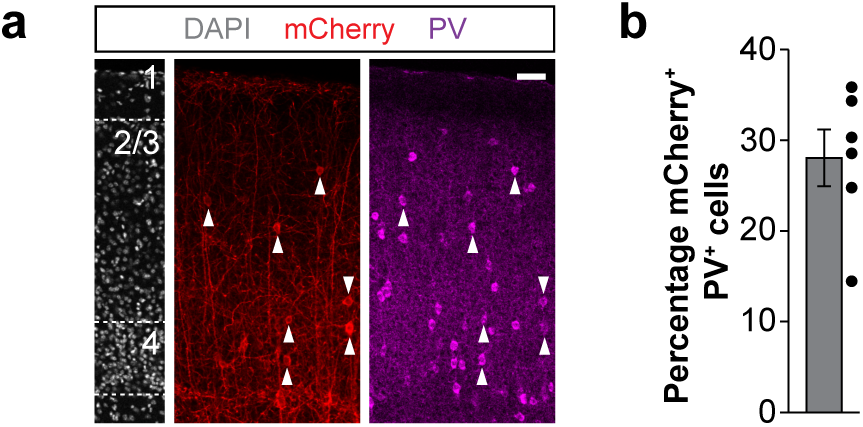
| Sparse infection of PV+ interneurons in L2/3 of S1. **a**, mCherry and PV expression in coronal sections through S1. **b**, quantification of the percentage of mCherry+ PV+ interneurons in L2/3 (*n* = 6 mice). Data are presented as mean ± s.e.m. Scale bar, 50 μm.

**Supplementary Fig. 2.**
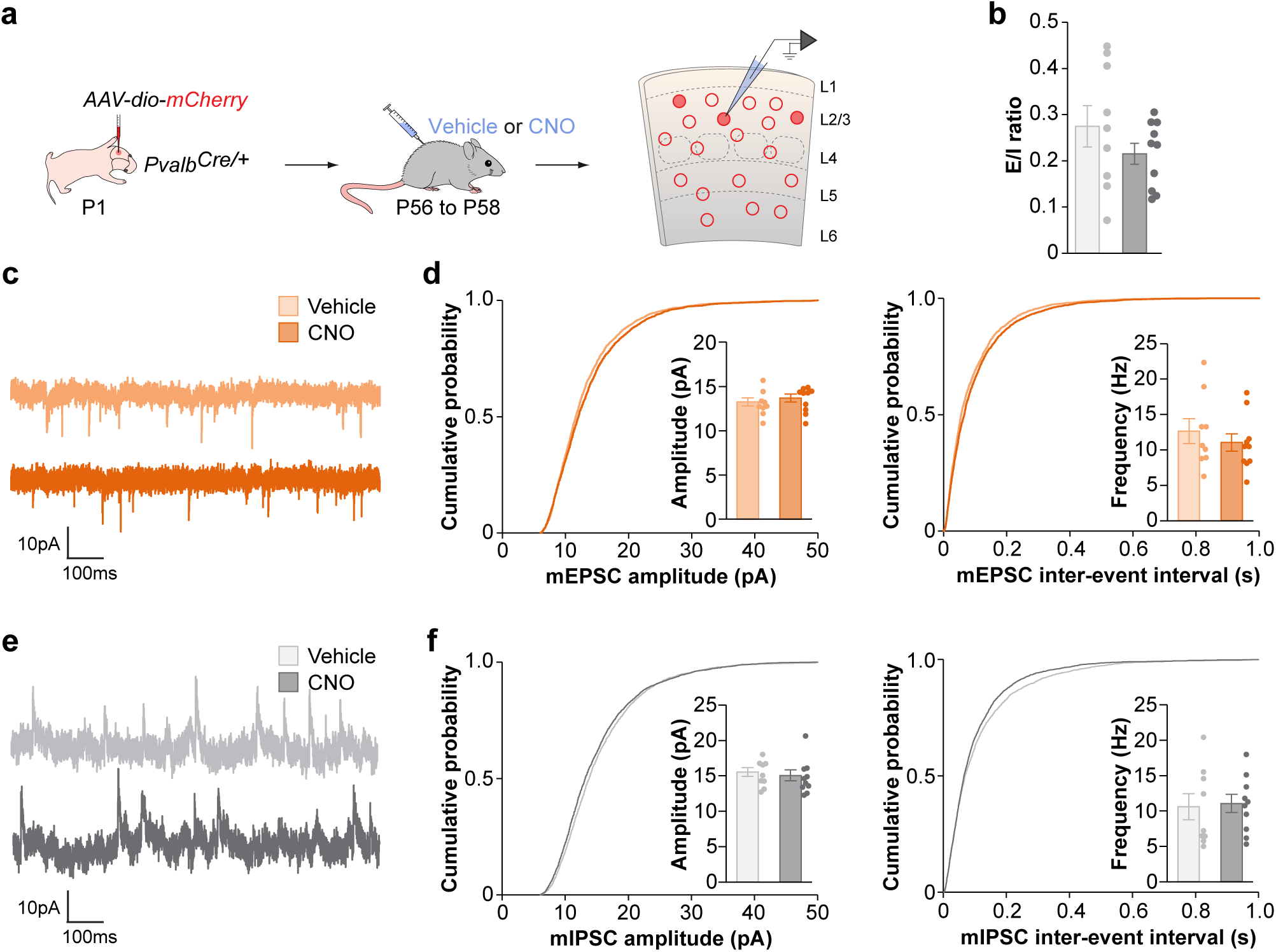
| CNO treatment does not affect the synaptic integration of mCherry-infected PV+ interneurons. **a**, Experimental strategy. **b**, Excitation/Inhibition (E/I) ratio in mCherry-infected PV+ interneurons in vehicle- and CNO-treated mice (vehicle, *n* = 9 cells; CNO, *n* = 10 cells; two-tailed Student’s t-test, *p* = 0.24). **c**, Traces of mEPSCs recorded from mCherry-infected PV+ interneurons in vehicle- and CNO-treated mice. **d**, Quantification of amplitude (vehicle, *n* = 9 cells; CNO, *n* = 10 cells; two-tailed Student’s t-test, *p* = 0.51) and frequency (vehicle, *n* = 9 cells; CNO, *n* = 10 cells; two-tailed Student’s t-test, *p* = 0.45) of mEPSCs recorded from mCherry-infected PV+ interneurons in vehicle- and CNO-treated mice. **e**, Traces of mIPSCs recorded from mCherry-infected PV+ interneurons in vehicle- and CNO-treated mice. **g**, Quantification of amplitude (vehicle, *n* = 9 cells; CNO, *n* = 10 cells; two-tailed Student’s t-test, *p* = 0.64) and frequency (vehicle, *n* = 9 cells; CNO, *n* = 10 cells; two-tailed Student’s t-test, *p* = 0.84) of mIPSCs recorded from mCherry-infected PV+ interneurons in vehicle- and CNO-treated mice. Data are presented as mean ± s.e.m.

**Supplementary Fig. 3.**
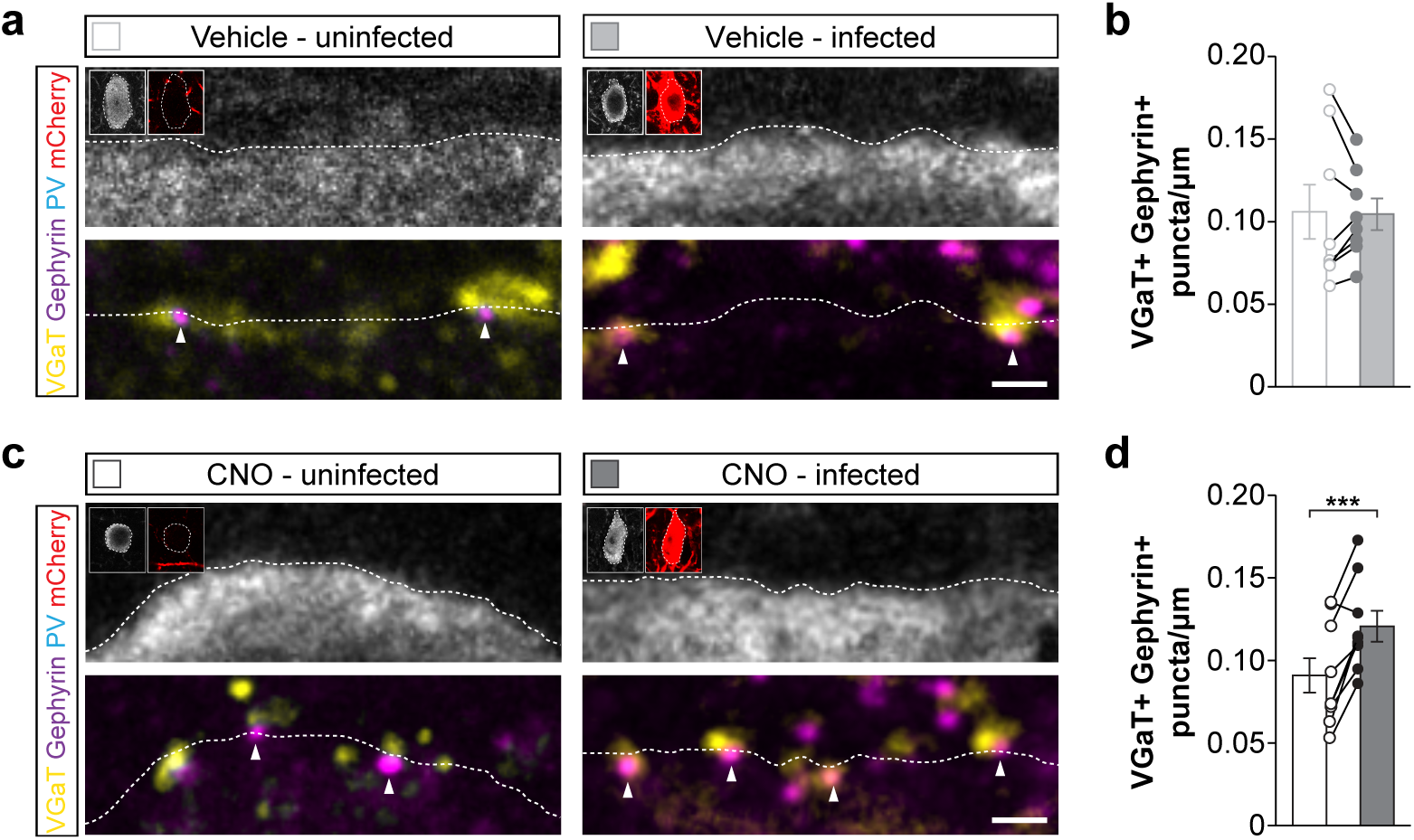
| Increasing activity of PV+ interneurons leads to an increased density of inhibitory synapses. **a**, Presynaptic VGaT+ puncta and postsynaptic Gephyrin+ clusters in neighbouring uninfected and hM3Dq-mCherry+ PV+ interneurons in vehicle-treated mice. Inserts show PV and mCherry immunoreactivity in cell somas. **b**, Quantification of the density of VGaT+/Gephyrin+ puncta on neighbouring uninfected and hM3D+ PV+ interneurons in vehicle-treated mice (*n* = 8; two-tailed paired Student’s t-test, *p* = 0.86). **c**, Presynaptic VGaT+ puncta and postsynaptic Gephyrin+ clusters in neighbouring uninfected and hM3Dq-mCherry+ PV+ interneurons in CNO-treated mice. Inserts show PV and mCherry immunoreactivity in the cell somas. **b,** Quantification of the density of VGaT+/Gephyrin+ puncta on neighbouring uninfected and hM3Dq-mCherry+ PV+ interneurons in CNO-treated mice (*n* = 9; two-tailed paired Student’s t-test, *p* < 0.001). Data are presented as mean ± s.e.m. Scale bars, 1 μm.

**Supplementary Fig. 4.**
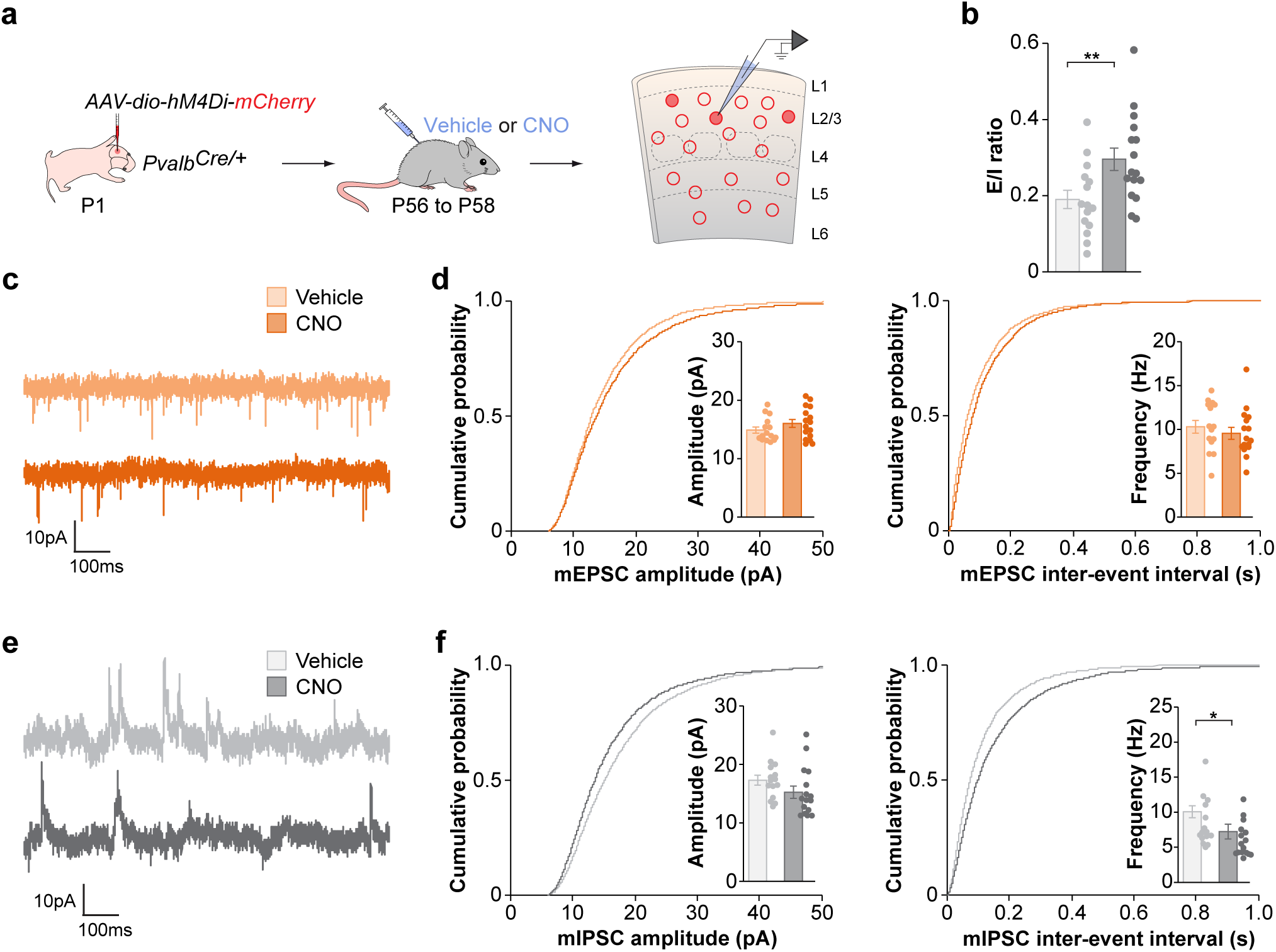
Reducing the activity of PV+ interneurons leads to reduced miniature inhibitory input. **a**, Experimental strategy. **b**, Excitation/Inhibition (E/I) ratio in hM4Di-mCherry infected PV interneurons in vehicle- and CNO-treated mice (vehicle, *n* = 15 cells; CNO, *n* = 16 cells; two-tailed Student’s t-test, *p* = 0.009). **c**, Traces of mEPSCs recorded from hM4Di-mCherry-infected PV+ interneurons in vehicle- and CNO-treated mice. **d**, Quantification of amplitude (vehicle, *n* = 15 cells; CNO, *n* = 16 cells; two-tailed Student’s t-test, *p* = 0.20) and frequency (vehicle, *n* = 15 cells; CNO, *n* = 16 cells; two-tailed Student’s t-test, *p* = 0.46) of mEPSCs recorded from hM3Dq-mCherry-infected PV+ interneurons in vehicle- and CNO-treated mice. **e**, Traces of mIPSCs recorded from hM4Di-mCherry-infected PV+ interneurons in vehicle- and CNO-treated mice. **f**, Quantification of amplitude (vehicle, *n* = 15 cells; CNO, *n* = 16 cells; two-tailed Student’s t-test, *p* = 0.14) and frequency (vehicle, *n* = 9 cells; CNO, *n* = 10 cells; two-tailed Student’s t-test, *p* = 0.03) of mIPSCs recorded from hM3Dq-mCherry-infected PV+ interneurons in vehicle- and CNO-treated mice. Data are presented as mean ± s.e.m.

**Supplementary Fig. 5.**
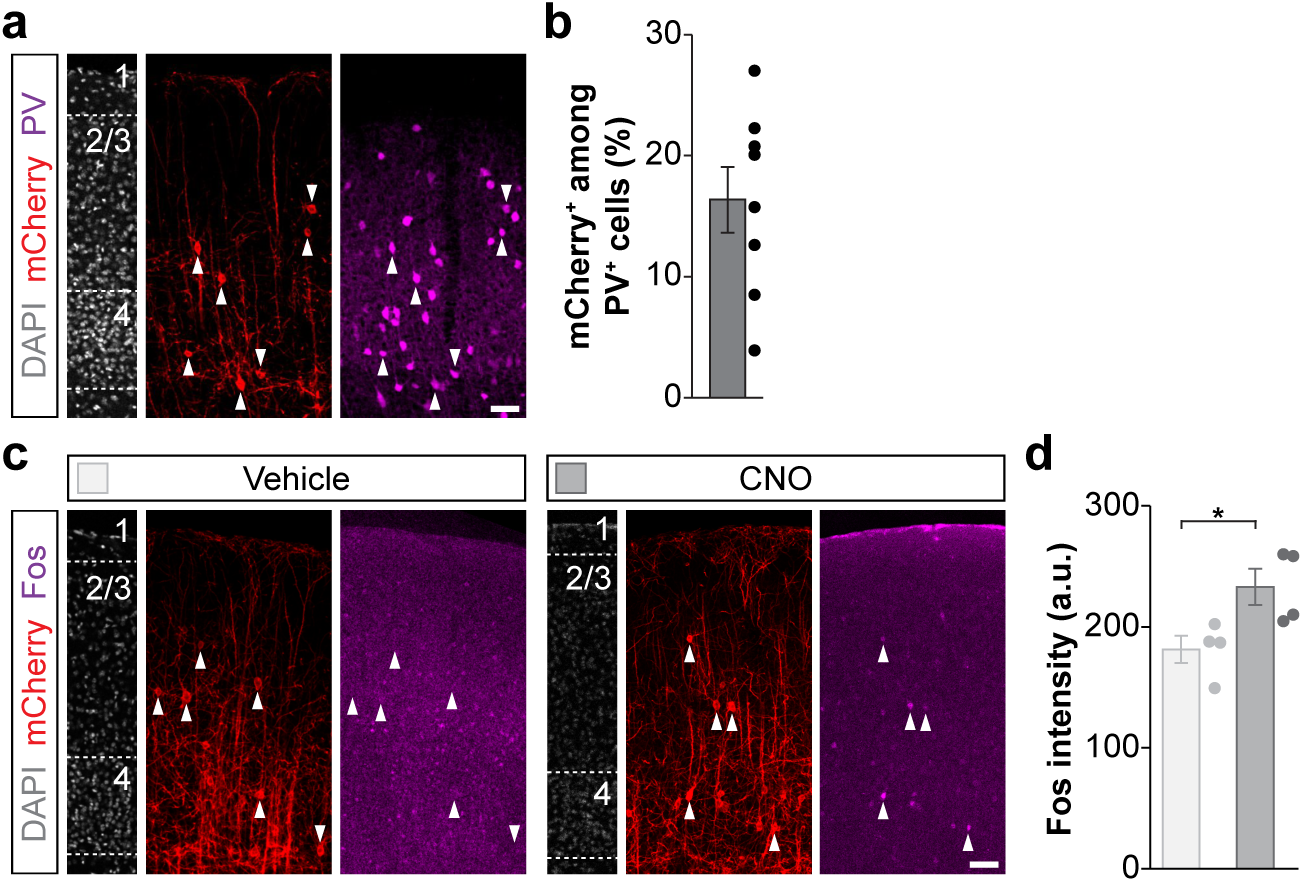
| Validation of Flp-dependent hM3Dq-mCherry AAV. **a**, PV and mCherry expression in coronal sections through S1. **b**, Quantification of the percentage of mCherry+ PV+ interneurons in L2/3 (*n* = 8 mice). **c,** Fos and mCherry expression in coronal sections through S1 from vehicle- and CNO-treated mice. **d,** quantification of the intensity of Fos staining in hM3Dq+ PV+ interneurons in vehicle- and CNO-treated mice (vehicle, *n* = 4 mice; CNO, *n* = 4 mice; two-tailed Student’s t-test, *p* = 0.03). Scale bars, 50 μm. Data are presented as mean ± s.e.m.

**Supplementary Fig. 6.**
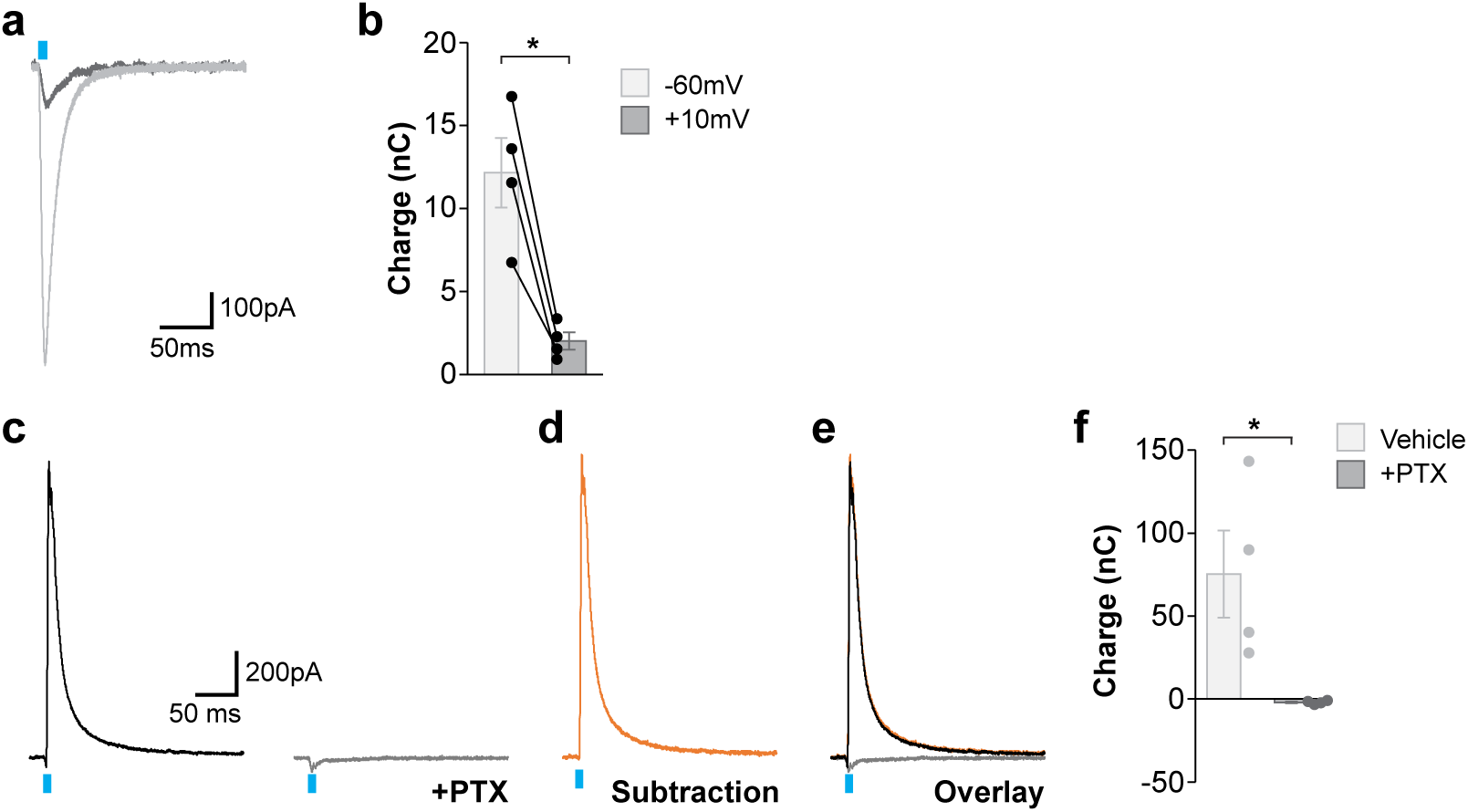
| Expression of channelrhodopsin in recorded cells does not interfere with eIPSC recordings. **a**, Traces of Channelrhodopsin (ChR2) evoked currents recorded from hM3Dq-mCherry expressing PV+ interneurons at a holding voltage of -60mV and +10mV in the presence of picrotoxin (PTX) at 100% LED power. **b**, Quantification of ChR2-evoked charge in the presence of PTX (−60mV, *n* = 4 cells; +10mV, *n* = 4 cells; two-tailed paired Student’s t-test, *p* = 0.01). **c,** Traces of eIPSCs recorded from an hM3Dq+ PV+ interneuron at a holding voltage of +10mV following stimulation of ChR2 expressing PV+ interneurons at 100% LED power in the absence or presence of picrotoxin (PTX). The PTX condition reflects the contribution of direct ChR2-evoked current (compare to panel a). **d,** Resultant trace of the subtraction of the traces shown in panel (c). **e,** Overlay of panels (c) and (d). **f,** Quantification of eIPSC charges recorded from hM3Dq+ PV+ interneurons following stimulation of ChR2+ PV+ interneurons at 100% LED power in the absence or presence of PTX (control, *n* = 4 cells; PTX, *n* = 4 cells; two-tailed Student’s t-test, *p* = 0.03). Blue boxes indicate LED stimulation. Data are presented as mean ± s.e.m.

**Supplementary Fig. 7.**
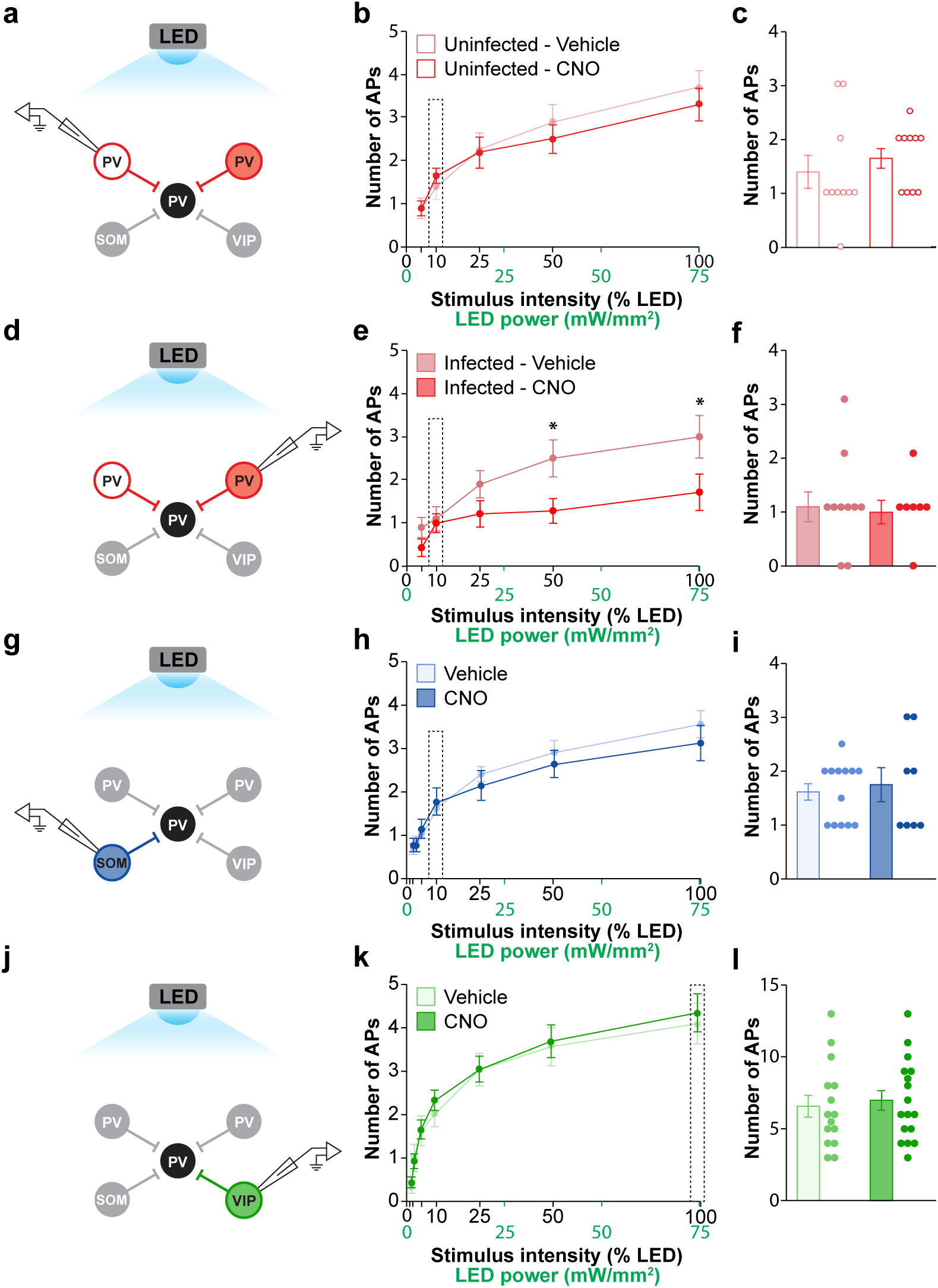
| Excitability of channelrhodopsin-expressing interneurons following LED stimulation at different intensities. **a**, Experimental strategy to record the excitability of uninfected PV+ interneurons in a cell-attached configuration in vehicle- and CNO-treated *Pvalb^Cre/Flp^;RCL^Chr2/+^* mice. **b**, Quantification of the number of elicited action potentials (APs) following LED stimulation at different intensities in uninfected PV+ interneurons from vehicle- and CNO-treated mice (5% LED-intensity, vehicle, *n* = 10 cells; CNO, *n* = 11 cells; two-tailed Student’s t-test, *p* = 0.97; 10% LED-intensity, vehicle, *n* = 10 cells; CNO, *n* = 11 cells; two-tailed Student’s t-test, *p* = 0.59; 25% LED-intensity, vehicle, *n* = 10 cells; CNO, *n* = 11 cells; two-tailed Student’s t-test, *p* = 0.76; 50% LED-intensity, vehicle, *n* = 10 cells; CNO, *n* = 11 cells; two-tailed Student’s t-test, *p* = 0.32; 100% LED-intensity, vehicle, *n* = 10 cells; CNO, *n* = 11 cells; two-tailed Student’s t-test, *p* = 0.29). **c**, Quantification of the selected stimulation intensity from panel (b). **d**, Experimental strategy to record the excitability of infected PV+ interneurons in a cell-attached configuration in vehicle- and CNO-treated *Pvalb^Cre/Flp^;RCL^Chr2/+^* mice. **e**, Quantification of the number of elicited APs following LED stimulation at different intensities in infected PV+ interneurons from vehicle- and CNO-treated mice (5% LED-intensity, vehicle, *n* = 10 cells; CNO, *n* = 10 cells; two-tailed Student’s t-test, *p* = 0.23; 10% LED-intensity, vehicle, *n* = 10 cells; CNO, *n* = 10 cells; two-tailed Student’s t-test, *p* = 1.00; 25% LED-intensity, vehicle, *n* = 10 cells; CNO, *n* = 10 cells; two-tailed Student’s t-test, *p* = 0.21; 50% LED-intensity, vehicle, *n* = 10 cells; CNO, *n* = 10 cells; two-tailed Student’s t-test, *p* = 0.04; 100% LED-intensity, vehicle, *n* = 10 cells; CNO, *n* = 10 cells; two-tailed Student’s t-test, *p* = 0.04). **f**, Quantification of the selected stimulation intensity from panel (e). **g**, Experimental strategy to record the excitability of SST+ interneurons in a cell-attached configuration in vehicle- and CNO-treated *Sst^Cre/+^;Pvalb^Flp/+^;RCL^Chr2/+^* mice. **h**, Quantification of the number of elicited APs following LED stimulation at different intensities in SOM+ interneurons in vehicle- and CNO-treated mice (2% LED-intensity, vehicle, *n* = 13 cells; CNO, *n* = 8 cells; two-tailed Student’s t-test, *p* = 0.65; 3% LED-intensity, vehicle, *n* = 13 cells; CNO, *n* = 8 cells; two-tailed Student’s t-test, *p* = 0.61; 5% LED-intensity, vehicle, *n* = 13 cells; CNO, *n* = 8 cells; two-tailed Student’s t-test, *p* = 0.59; 10% LED-intensity, vehicle, *n* = 13 cells; CNO, *n* = 8 cells; two-tailed Student’s t-test, *p* = 0.67; 25% LED-intensity, vehicle, *n* = 13 cells; CNO, *n* = 8 cells; two-tailed Student’s t-test, *p* = 0.48; 50% LED-intensity, vehicle, *n* = 13 cells; CNO, *n* = 8 cells; two-tailed Student’s t-test, *p* = 0.55; 100% LED-intensity, vehicle, *n* = 13 cells; CNO, *n* = 8 cells; two-tailed Student’s t-test, *p* = 0.42). **i**, Quantification of the selected stimulation intensity from panel (h). **j**, Experimental strategy to record the excitability of VIP+ interneurons in a cell-attached configuration in vehicle- and CNO-treated *Vip^Cre/+^;Pvalb^Flp/+^;RCL^Chr2/+^* mice. **k**, Quantification of the number of elicited APs following LED stimulation at different intensities in VIP+ interneurons in vehicle- and CNO-treated mice (2% LED-intensity, vehicle, *n* = 15 cells; CNO, *n* = 17 cells; two-tailed Student’s t-test, *p* = 0.56; 3% LED-intensity, vehicle, *n* = 15 cells; CNO, *n* = 17 cells; two-tailed Student’s t-test, *p* = 0.77; 5% LED-intensity, vehicle, *n* = 15 cells; CNO, *n* = 17 cells; two-tailed Student’s t-test, *p* = 0.94; 10% LED-intensity, vehicle, *n* = 15 cells; CNO, *n* = 17 cells; two-tailed Student’s t-test, *p* = 0.38; 25% LED-intensity, vehicle, *n* = 15 cells; CNO, *n* = 17 cells; two-tailed Student’s t-test, *p* = 0.98; 50% LED-intensity, vehicle, *n* = 15 cells; CNO, *n* = 17 cells; two-tailed Student’s t-test, *p* = 0.85; 100% LED-intensity, vehicle, *n* = 15 cells; CNO, *n* = 17 cells; two-tailed Student’s t-test, *p* = 0.69). **l**, Quantification of the selected stimulation intensity from panel (k). Data are presented as mean ± s.e.m. Dotted boxes indicate the selected stimulation strength.

**Supplementary Fig. 8.**
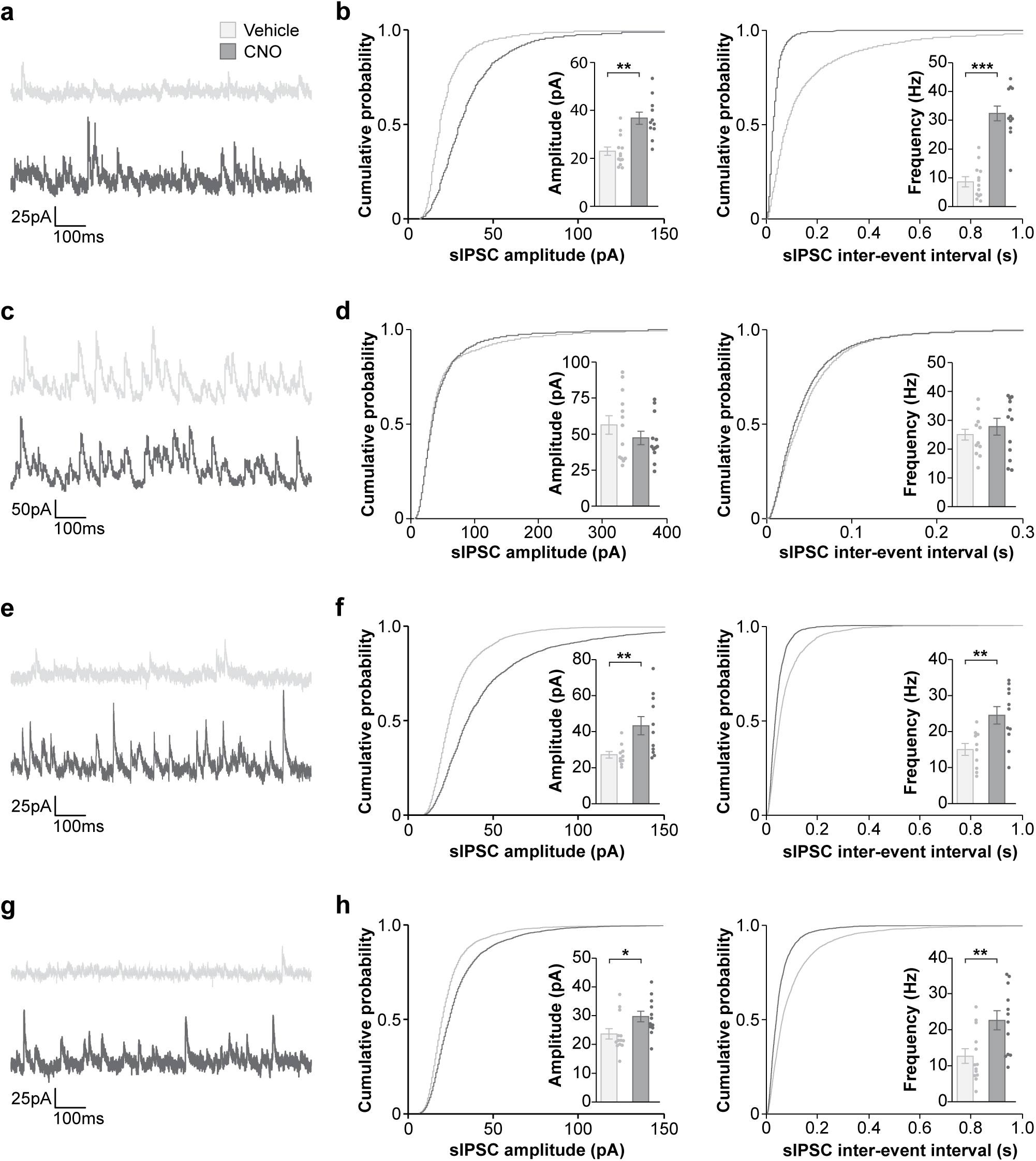
| Spontaneous IPSC recordings confirm the increase in PV+ interneuron activity. **a**, Traces of spontaneous IPSCs (sIPSCs) recorded from hM3Dq+ PV+ interneurons in vehicle- and CNO-treated *Pvalb^Cre/Flp^;RCL^Chr2/+^* mice. **b**, Quantification of amplitude (vehicle, *n* = 16 cells; CNO, *n* = 13 cells; two-tailed Student’s t-test, *p* = 0.003) and frequency (vehicle, *n* = 16 cells; CNO, *n* = 13 cells; two-tailed Student’s t-test, *p* < 0.001) of sIPSCs recorded from hM3Dq+ PV+ interneurons in vehicle- and CNO-treated *Pvalb^Cre/Flp^;RCL^Chr2/+^* mice. **c**, Traces of sIPSCs recorded from L2/3 pyramidal neurons in vehicle- and CNO-treated *Pvalb^Cre/Flp^;RCL^Chr2/+^* mice. **d**, Quantification of amplitude (vehicle, *n* = 13 cells; CNO, *n* = 12 cells; two-tailed Student’s t-test, *p* = 0.28) and frequency (vehicle, *n* = 13 cells; CNO, *n* = 12 cells; two-tailed Student’s t-test, *p* = 0.43) of sIPSCs recorded from L2/3 pyramidal neurons in vehicle- and CNO-treated *Pvalb^Cre/Flp^;RCL^Chr2/+^* mice. **e**, Traces of sIPSCs recorded from hM3Dq+ PV+ interneurons in vehicle- and CNO-treated *Sst^Cre/+^;Pvalb^Flp/+^;RCL^Chr2/+^* mice. **f**, Quantification of amplitude (vehicle, *n* = 10 cells; CNO, *n* = 11 cells; two-tailed Student’s t-test, *p* = 0.009) and frequency (vehicle, *n* = 10 cells; CNO, *n* = 11 cells; two-tailed Student’s t-test, *p* = 0.005) of sIPSCs recorded from hM3Dq+ PV+ interneurons in vehicle- and CNO-treated *Sst^Cre/+^;Pvalb^Flp/+^;RCL^Chr2/+^* mice. **g**, Traces of sIPSCs recorded from hM3Dq+ PV+ interneurons in vehicle- and CNO-treated *Vip^Cre/+^;Pvalb^Flp/+^;RCL^Chr2/+^* mice. **h**, Quantification of amplitude (vehicle, *n* = 13 cells; CNO, *n* = 12 cells; two-tailed Student’s t-test, *p* = 0.03) and frequency (vehicle, *n* = 13 cells; CNO, *n* = 12 cells; two-tailed Student’s t-test, *p* = 0.006) of sIPSCs recorded from hM3Dq+ PV+ interneurons in vehicle- and CNO-treated *Vip^Cre/+^;Pvalb^Flp/+^;RCL^Chr2/+^* mice. Data are presented as mean ± s.e.m.

**Supplementary Fig. 9.**
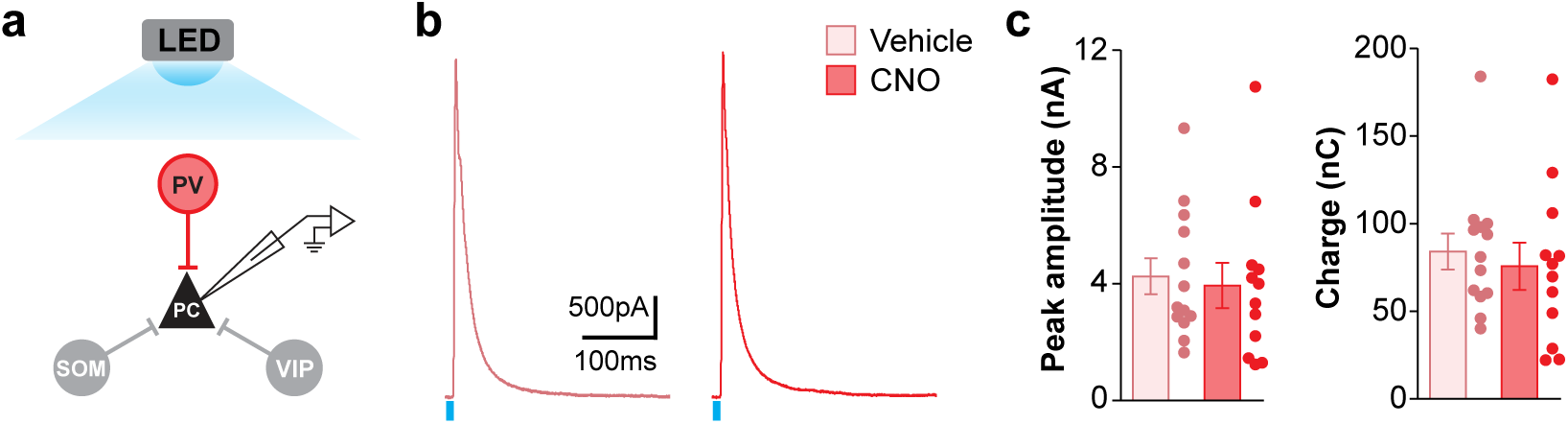
| Increased activity in PV+ interneurons does not increase PV+ input onto pyramidal neurons. **a**, Experimental strategy. **b**, Traces of eIPSCs recording from layer 2/3 pyramidal cells following full-field stimulation of PV+ interneurons in vehicle- and CNO-treated *Pvalb^Cre/Flp^;RCL^Chr2/+^* mice. **c**, Quantification of the peak amplitude (vehicle, *n* = 13 cells; CNO, *n* = 12 cells; two-tailed Student’s t-test, *p* = 0.76) and charge (vehicle, *n* = 13 cells; CNO, *n* = 12 cells; two-tailed Student’s t-test, *p* = 0.66) of eIPSCs evoked by LED stimulation in layer 2/3 pyramidal cells following full-field stimulation of PV+ interneurons in vehicle- and CNO-treated *Pvalb^Cre/Flp^;RCL^Chr2/+^* mice. Data are presented as mean ± s.e.m.

**Supplementary Fig. 10.**
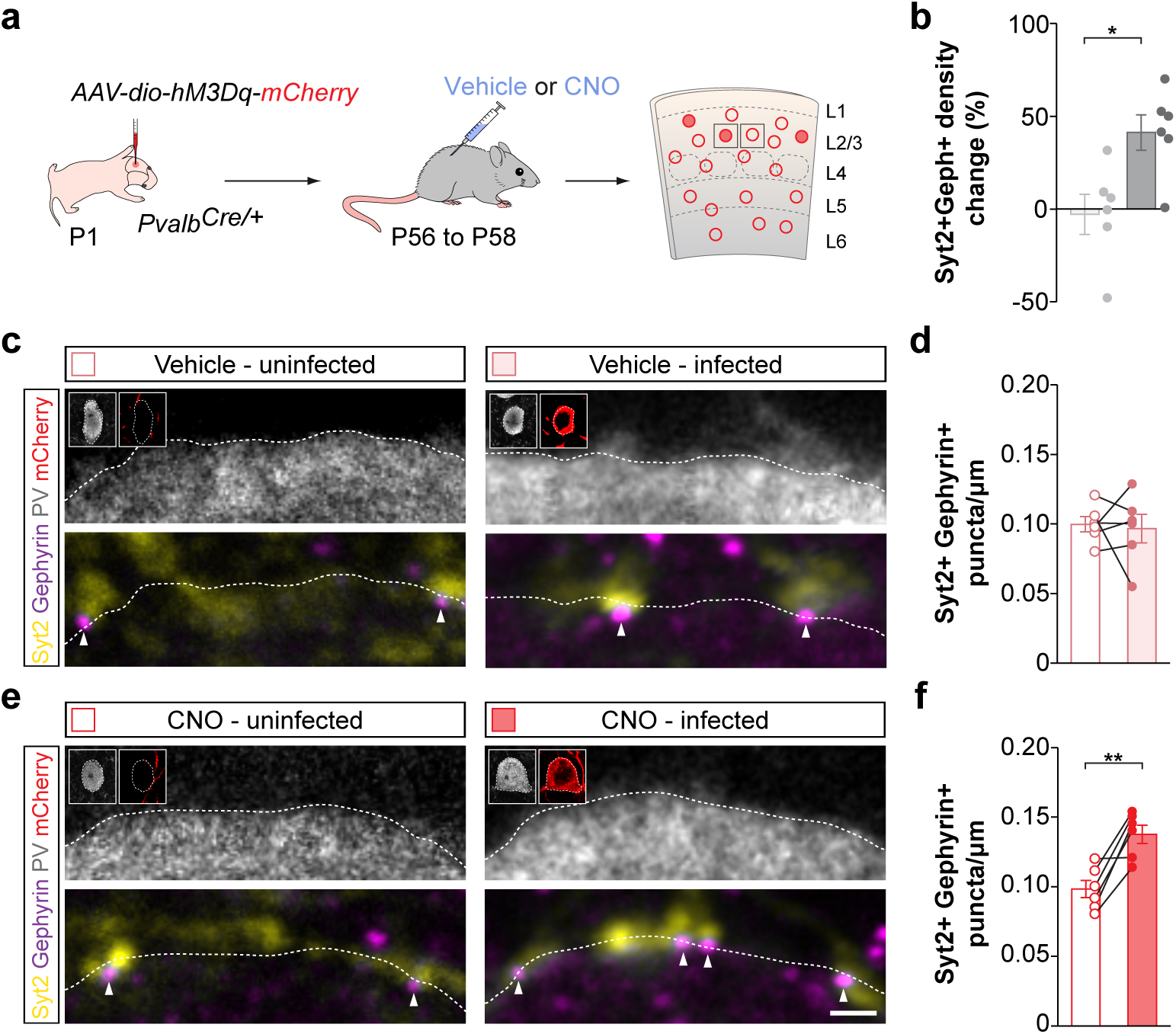
| Increasing the activity of PV+ interneurons leads to an increase in inhibitory synaptic density. **a**, Schematic outline of the experimental strategy. **b**, Quantification of change in synaptic density between infected and uninfected PV+ interneurons in vehicle- and CNO-treated mice (vehicle, *n* = 6 mice; CNO, *n* = 6 mice; two-tailed Student’s t-test, *p* = 0.01). **c,** Confocal images illustrating presynaptic Syt2+ puncta and postsynaptic Gephyrin+ clusters in neighbouring uninfected and hM3Dq-mCherry expressing PV+ interneurons in vehicle-treated mice. Inserts show PV and mCherry immunoreactivity of the cell’s soma. Scale bar 1 μm. **d,** Quantification of the density of colocalizing Syt2+/Gephyrin+ puncta on neighbouring uninfected and hM3Dq expressing PV+ interneurons in vehicle-treated mice (*n* = 6, ; two-tailed paired Student’s t-test, *p* = 0.79). **e,** Confocal images illustrating presynaptic Syt2+ puncta and postsynaptic Gephyrin+ clusters in neighbouring uninfected and hM3Dq-mCherry expressing PV+ interneurons in CNO-treated mice. Inserts show PV and mCherry immunoreactivity of the cell’s soma. Scale bar 1 μm. **d,** Quantification of the density of colocalizing Syt2+/Gephyrin+ puncta on neighbouring uninfected and hM3Dq expressing PV+ interneurons in CNO-treated mice (*n* = 6, ; two-tailed paired Student’s t-test, *p* = 0.006). Data are presented as mean ± s.e.m.

**Supplementary Fig. 11.**
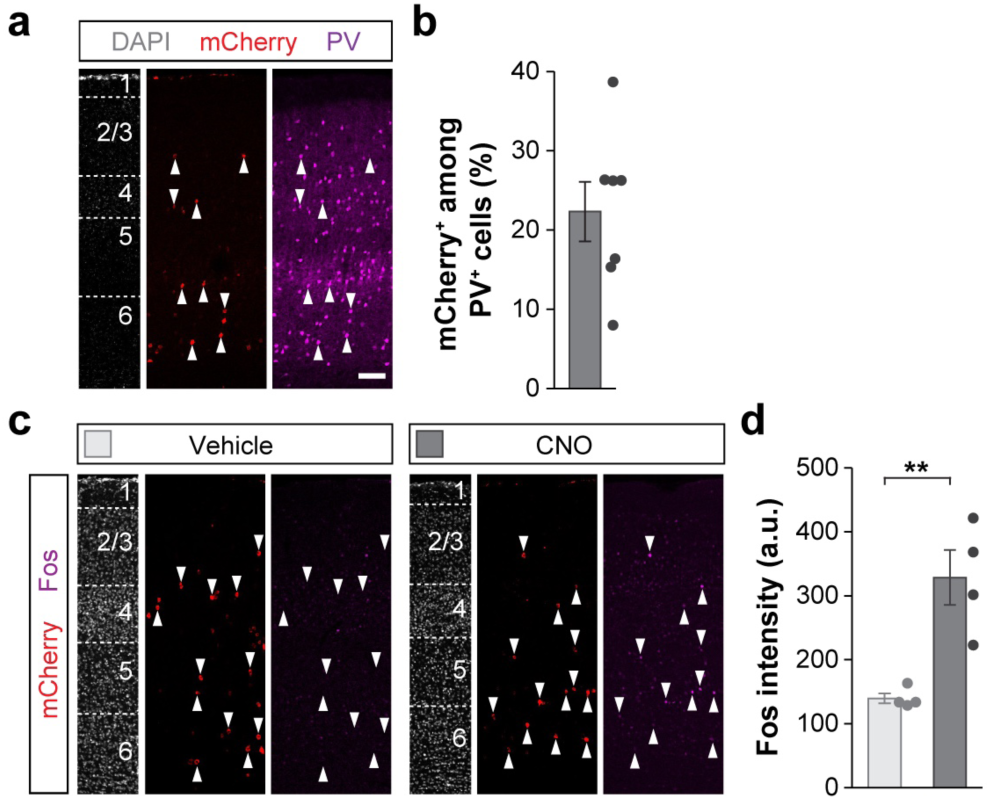
| Validation of *HA-vTRAP-hM3Dq-myc* AAV. **a**, PV and mCherry expression in coronal sections through S1. **b**, Quantification of the percentage of mCherry+ PV+ interneurons in L2/3 (*n* = 7 mice). **c,** Fos and mCherry expression in coronal sections through S1 from vehicle- and CNO-treated mice. **d,** Quantification of the intensity of Fos staining in HA-vTRAP-hM3Dq-myc infected PV+ interneurons in vehicle- and CNO-treated mice (vehicle, *n* = 4 mice; CNO, *n* = 4 mice; two-tailed Student’s t-test, *p* = 0.004). Scale bar, 50 μm. Data are presented as mean ± s.e.m.

**Supplementary Fig. 12.**
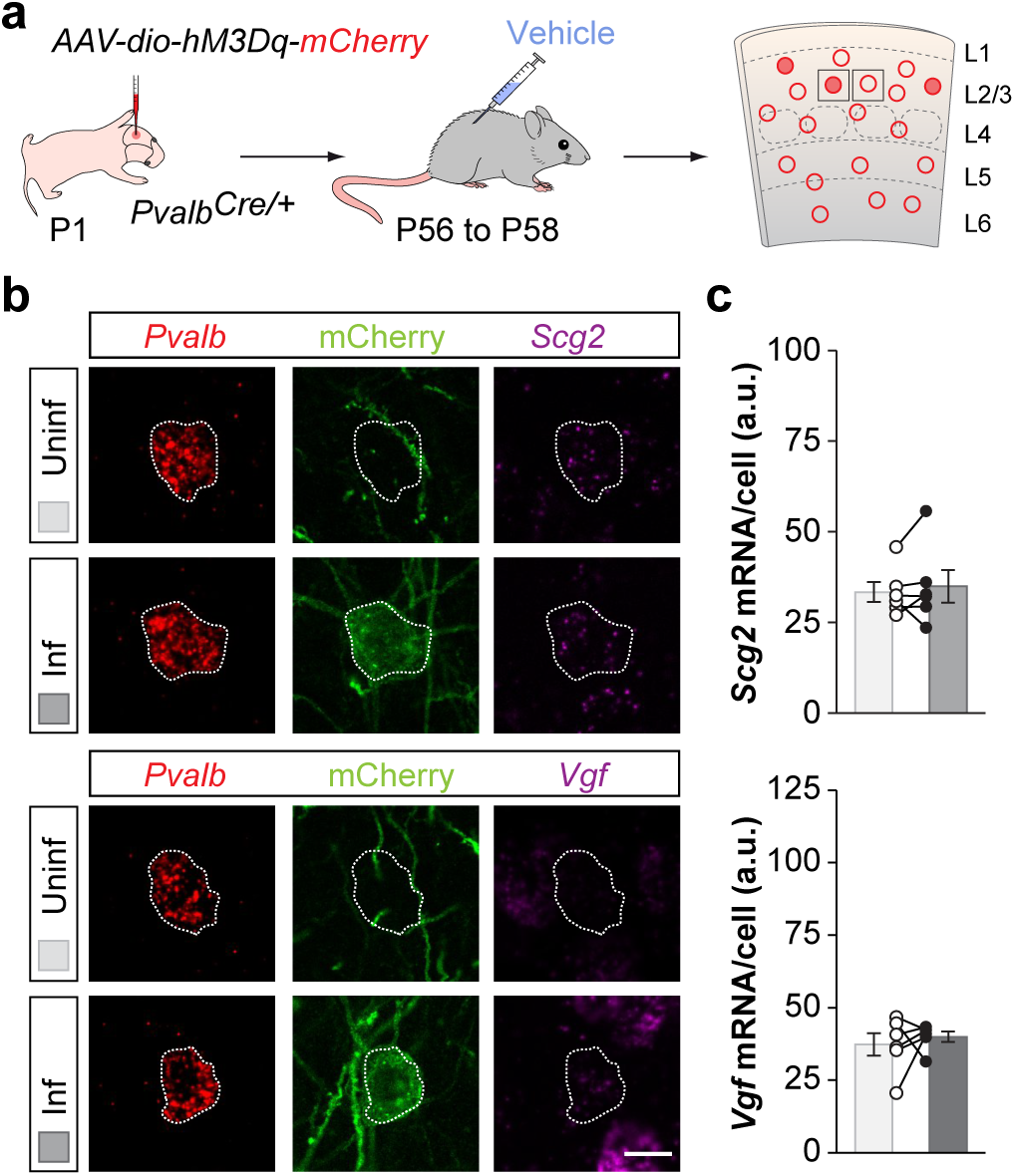
| Vehicle treatment of hM3Dq+ PV+ interneurons does not affect *Scg2* and *Vgf* expression. **a**, Experimental strategy. **b**, *Scg2* and *Vgf* mRNA expression in neighbouring infected and uninfected PV+ interneurons in vehicle-treated mice (*n* = 6 mice; two-tailed paired Student’s t-test, *p* = 0.54). **c**, Quantification of expression of *Scg2* and *Vgf* in neighbouring infected and uninfected PV+ interneurons in vehicle-treated mice (*n* = 5 mice; two-tailed paired Student’s t-test, *p* = 0.58). Data are presented as mean ± s.e.m. Scale bar, 10 μm.

**Supplementary Fig. 13.**
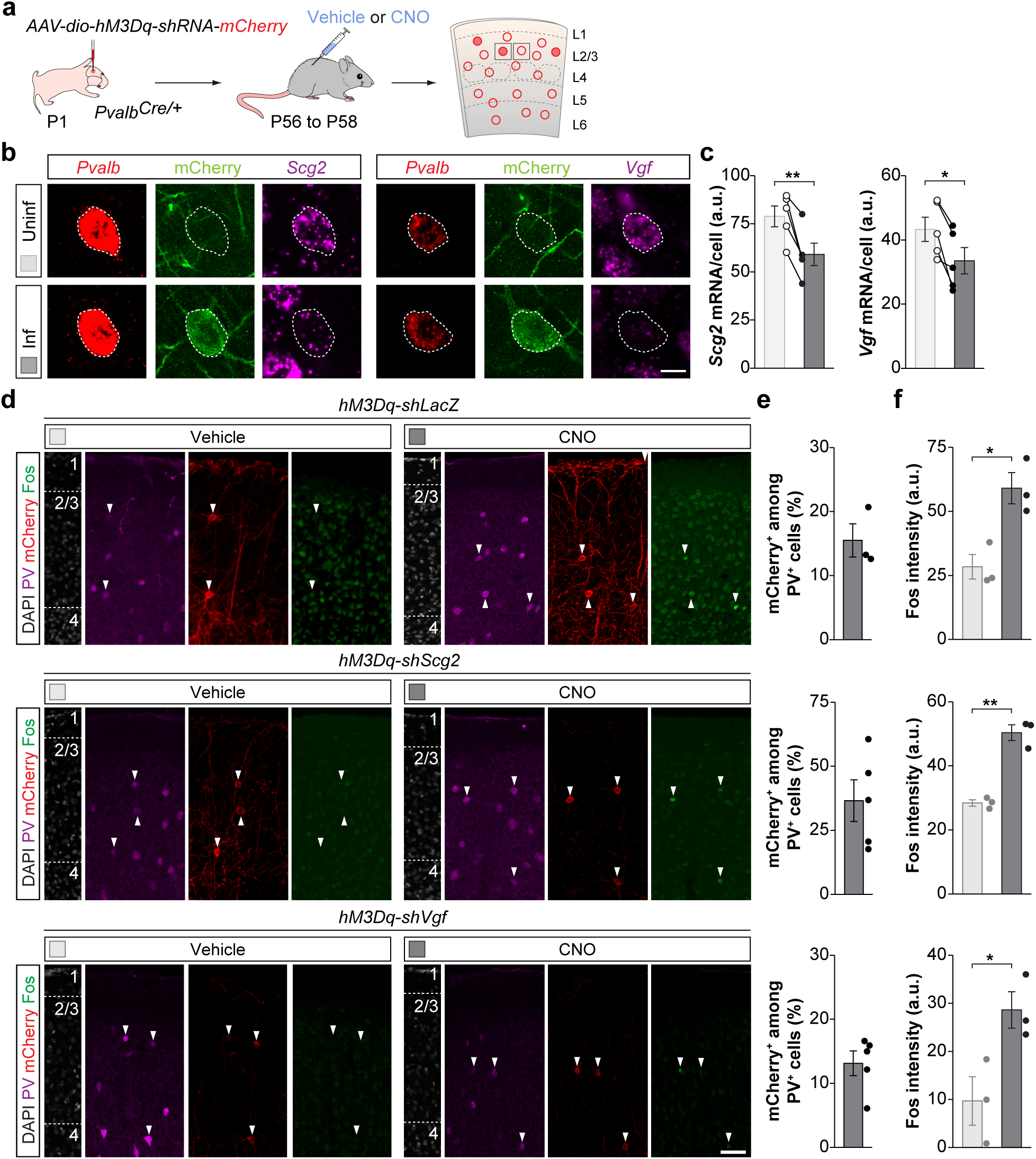
| Validation of shRNA AAVs. **a**, Experimental strategy. **b**, *Scg2* and *Vgf* mRNA expression in neighbouring infected and uninfected PV+ interneurons in CNO-treated mice. **c**, Quantification of *Scg2* and *Vgf* mRNAs (*shScg2-mCherry*: *n* = 6 mice; two-tailed paired Student’s t-test, *p* = 0.007; *shVgf-mCherry*: *n* = 6 mice; two-tailed paired Student’s t-test, *p* = 0.02). **d**, Fos, mCherry and PV expression in coronal sections through S1 from vehicle- and CNO-treated mice infected with *LacZ-mCherry*, *shScg2-mCherry*, and *Vgf-mCherry*. **e**, Quantification of the percentage of mCherry+ PV+ interneurons in L2/3 of mice infected with *LacZ-mCherry* (*n* = 3 mice), *shScg2-mCherry* (*n* = 5 mice), and *Vgf-mCherry* (*n* = 5 mice). **f**, Quantification of the intensity of Fos staining in PV+ interneurons in vehicle- and CNO-treated mice infected with *LacZ-mCherry*, *shScg2-mCherry*, and *Vgf-mCherry* (*shLacZ-mCherry*: vehicle, *n* = 3 mice; CNO, *n* = 3 mice; two-tailed Student’s t-test, *p* = 0.02; *shScg2-mCherry*: vehicle, *n* = 3 mice; CNO, *n* = 3 mice; two-tailed Student’s t-test, *p* = 0.001; *Vgf-mCherry*: vehicle, *n* = 3 mice; CNO, *n* = 3 mice; two-tailed Student’s t-test, *p* = 0.04). Data are presented as mean ± s.e.m. Scale bars, 10 μm (b) and 50 μm (d).

**Supplementary Fig. 14.**
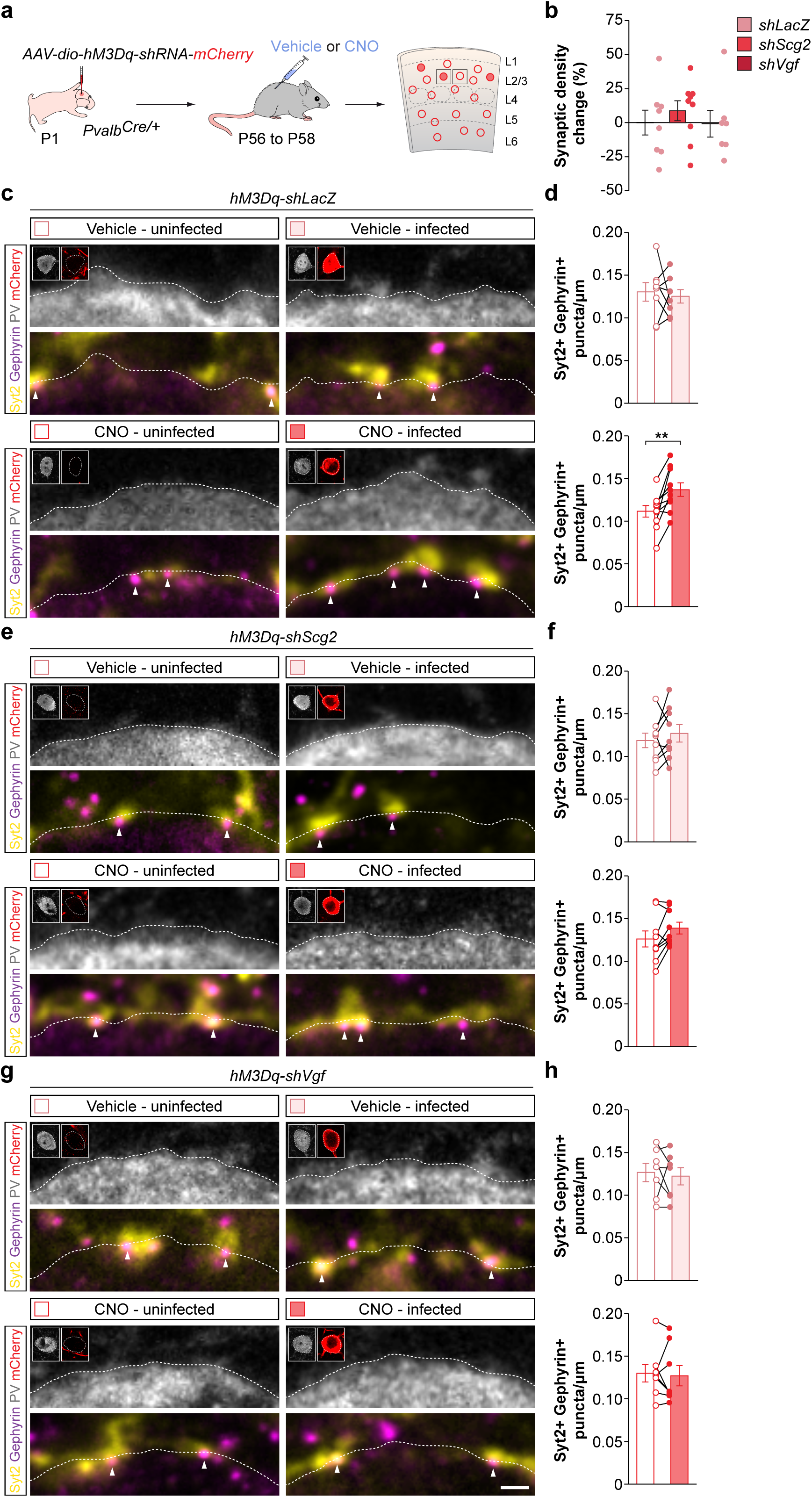
| Downregulation of *Scg2* and *Vgf* prevents the increase in synaptic density induced by increased activity. **a**, Experimental strategy. **b**, Quantification of the change in synaptic density between infected and uninfected PV+ interneurons in vehicle-treated mice injected with *hM3Dq-shLacZ* (*n* = 8 mice), *hM3Dq-shScg2* (*n* = 9 mice, one-sample t-test, *p* = 0.27) and *hM3Dq-shVgf* (*n* = 7 mice, one-sample t-test, *p* = 0.94). **c**, Presynaptic Syt2+ puncta and postsynaptic Gephyrin+ clusters in neighbouring uninfected and infected PV+ interneurons in vehicle- and CNO-treated mice injected with *shLacZ-mCherry*. Inserts show PV and mCherry immunoreactivity in the cell somas. **d,** Quantification of the density of colocalizing Syt2+/Gephyrin+ puncta on neighbouring uninfected and infected PV+ interneurons in vehicle- and CNO-treated mice injected with *shLacZ-mCherry* (vehicle: *n* = 8 ; two-tailed paired Student’s t-test, *p* = 0.67; CNO: *n* = 10; two-tailed paired Student’s t-test, *p* = 0.003). **e**, Presynaptic Syt2+ puncta and postsynaptic Gephyrin+ clusters in neighbouring uninfected and infected PV+ interneurons in vehicle- and CNO-treated mice injected with *shScg2-mCherry*. Inserts show PV and mCherry immunoreactivity in the cell somas. **f**, Quantification of the density of colocalizing Syt2+/Gephyrin+ puncta on neighbouring uninfected and infected PV+ interneurons in vehicle- and CNO-treated mice injected with *shScg2-mCherry* (vehicle: *n* = 9; two-tailed paired Student’s t-test, *p* = 0.40; CNO: *n* = 9 ; two-tailed paired Student’s t-test, *p* = 0.07). **g**, Presynaptic Syt2+ puncta and postsynaptic Gephyrin+ clusters in neighbouring uninfected and infected PV+ interneurons in vehicle- and CNO-treated mice injected with *shVgf-mCherry*. Inserts show PV and mCherry immunoreactivity in the cell somas. **h**, Quantification of the density of colocalizing Syt2+/Gephyrin+ puncta on neighbouring uninfected and infected PV+ interneurons in vehicle- and CNO-treated mice injected with *shVgf -mCherry* (vehicle: *n* = 7; two-tailed paired Student’s t-test, *p* = 0.70; CNO: *n* = 8; two-tailed paired Student’s t-test, *p* = 0.72). Data are presented as mean ± s.e.m. Scale bar, 1 μm.

**Supplementary Fig. 15.**
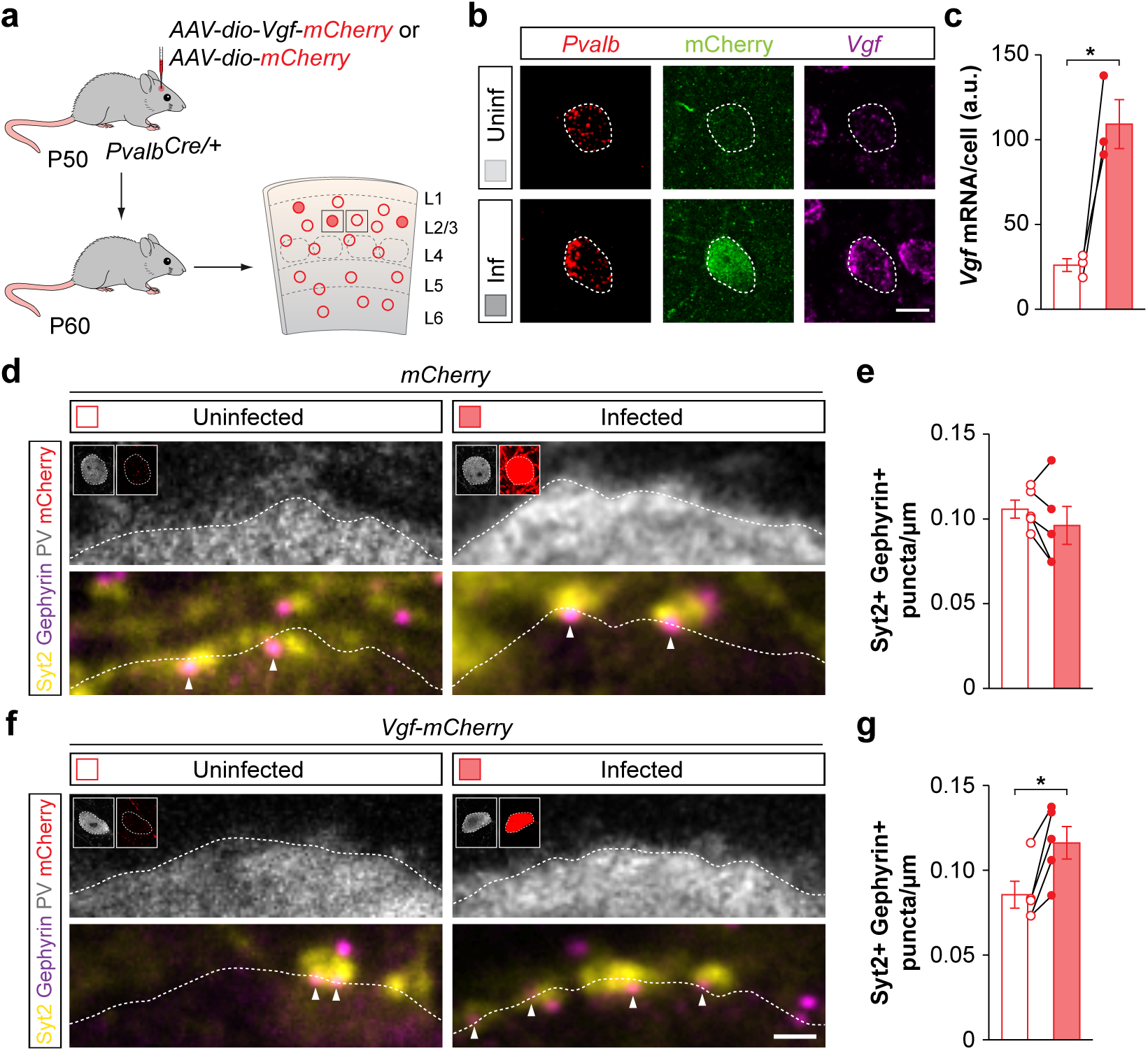
Overexpression of *Vgf* increases PV-PV interconnectivity. **a**, Experimental strategy. **b,** *Vgf* mRNA expression in neighbouring uninfected and Vgf-mCherry+ PV+ interneurons. **c**, Quantification of *Vgf* mRNA expression (*n* = 3 mice; two-tailed paired Student’s t-test, *p* = 0.03). **d**, Presynaptic Syt2+ puncta and postsynaptic Gephyrin+ clusters in neighbouring uninfected and infected mCherry+ PV+ interneurons. Inserts show PV and mCherry immunoreactivity in the cell somas. **e,** Quantification of the density of colocalizing Syt2+/Gephyrin+ puncta on neighbouring uninfected and mCherry+ PV+ interneurons (*n* = 5; two-tailed paired Student’s t-test, *p* = 0.23). **f,** Presynaptic Syt2+ puncta and postsynaptic Gephyrin+ clusters in neighbouring uninfected and Vgf-mCherry+ PV+ interneurons. Inserts show PV and mCherry immunoreactivity in the cell somas. **g,** Quantification of the density of colocalizing Syt2+/Gephyrin+ puncta on neighbouring uninfected and Vgf-mCherry+ PV+ interneurons (*n* = 5; two-tailed paired Student’s t-test, *p* = 0.01).

**Supplementary Table 1.**
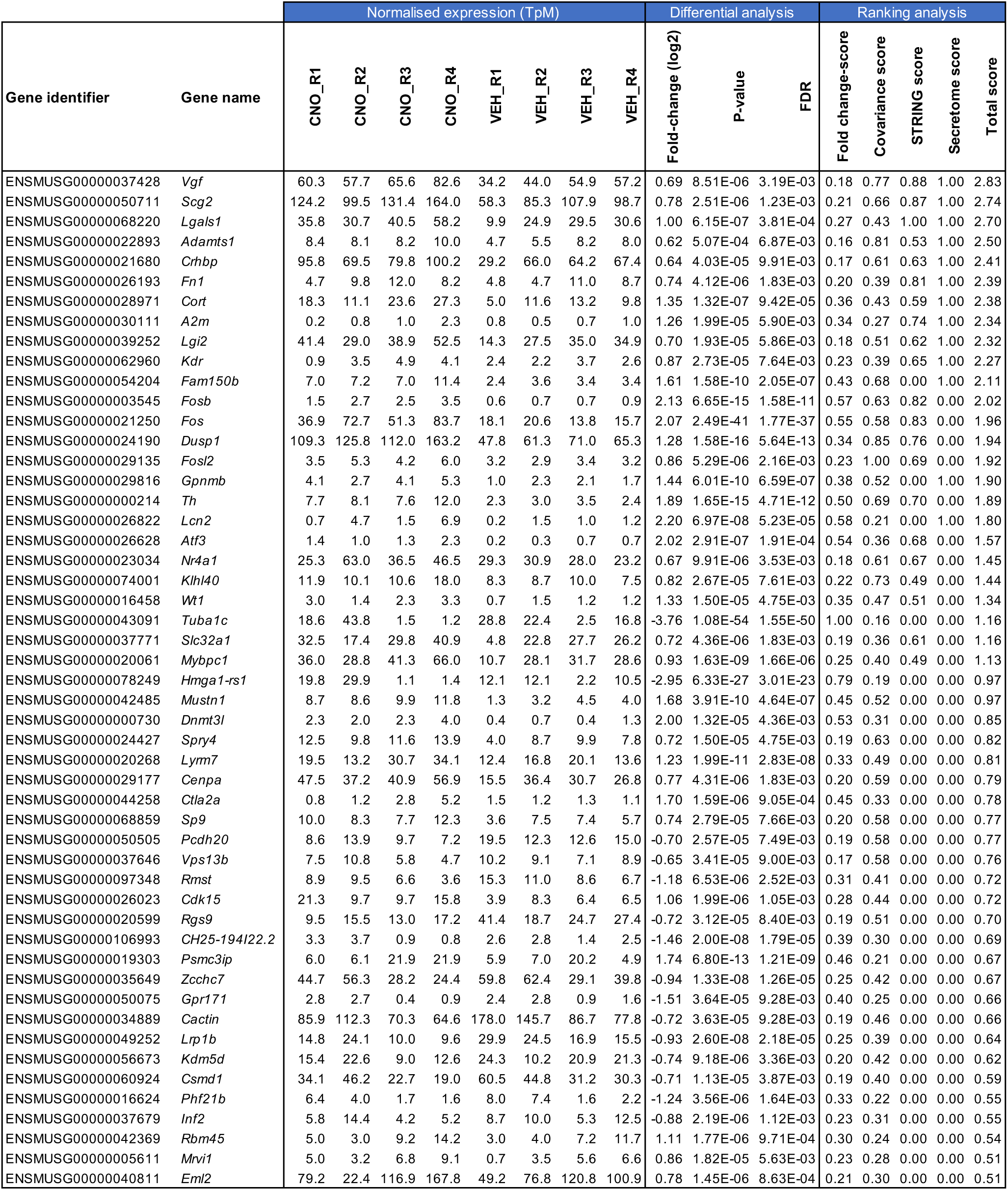
Genes up and downregulated following PV+ interneuron activation

**Supplementary Table 2.**
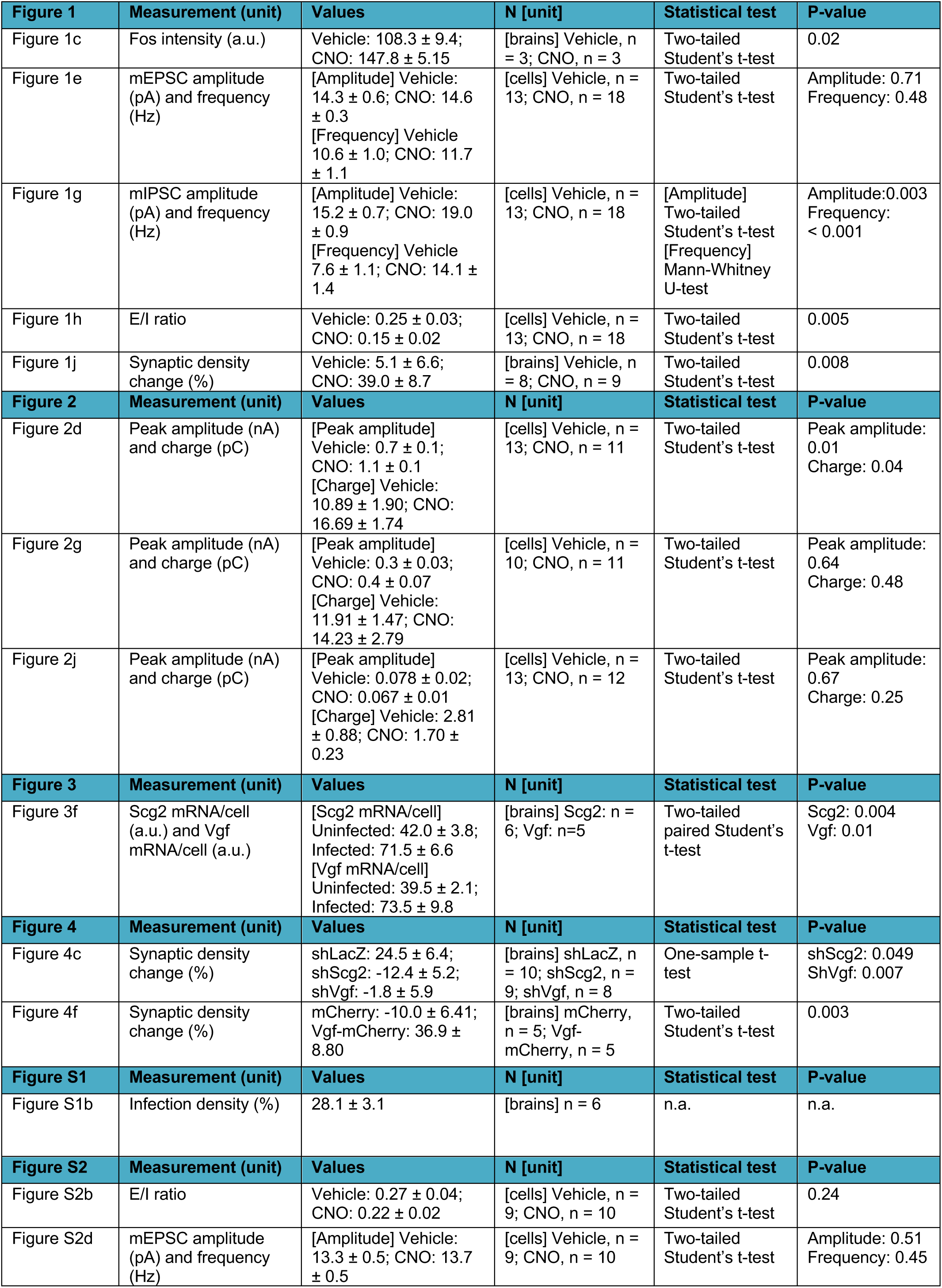
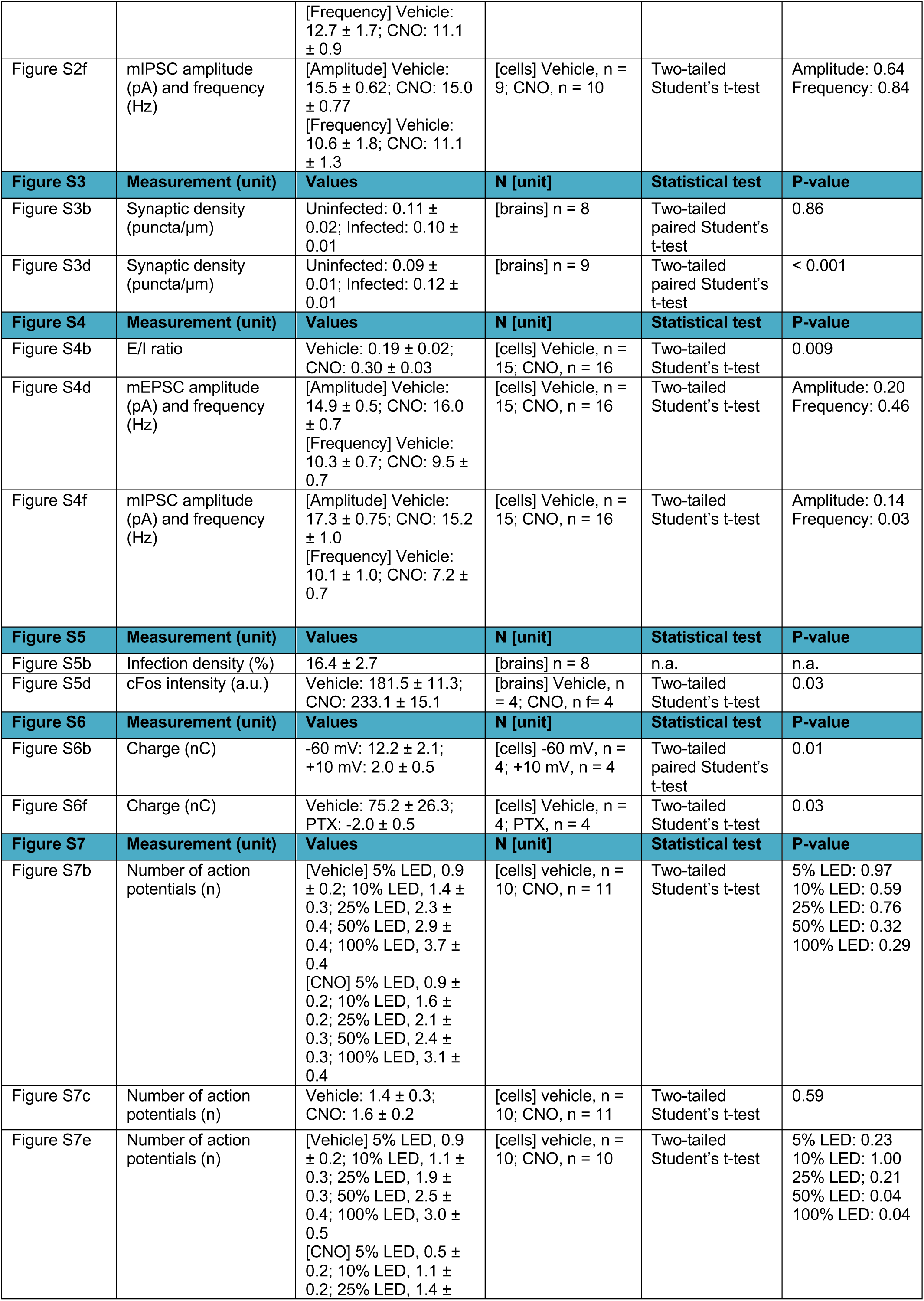
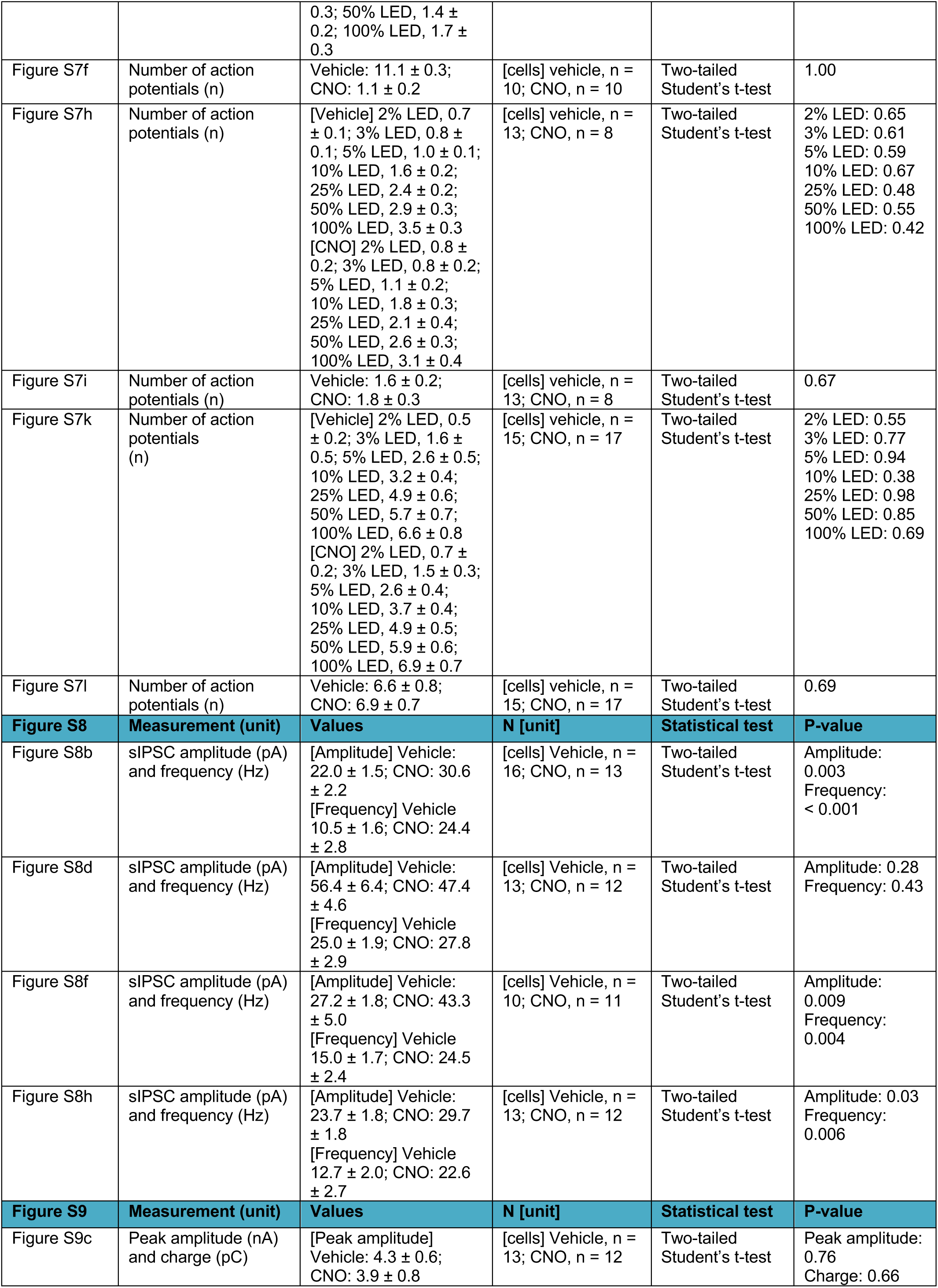
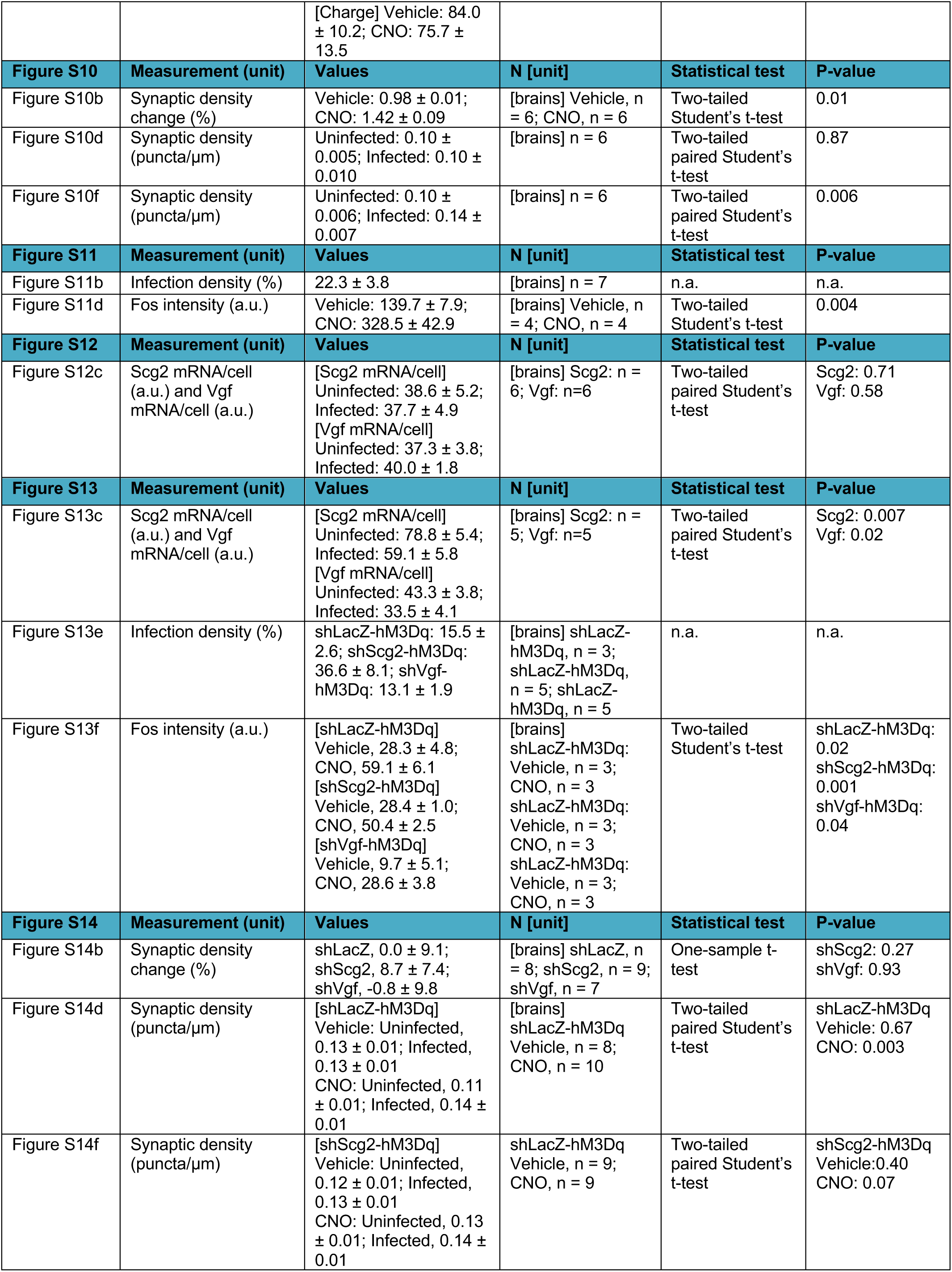
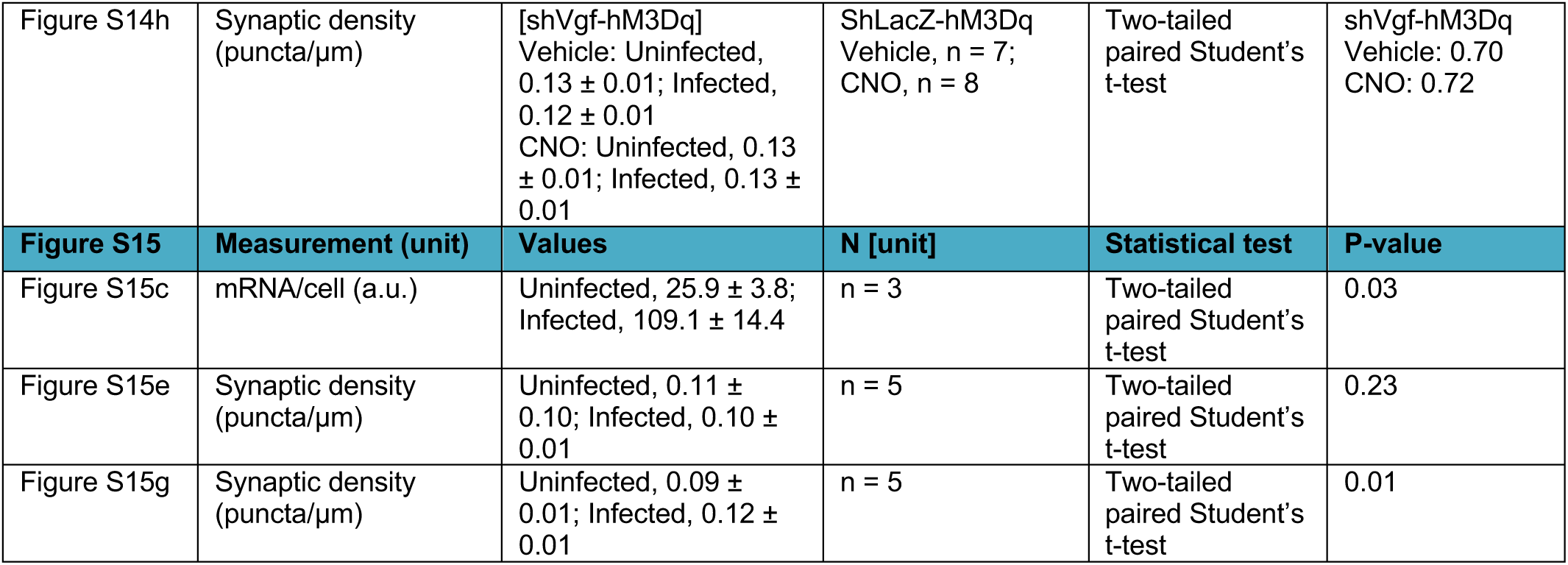
Summary of data and statistical analyses.

## Notes

### Competing Interest Statement

The authors have declared no competing interest.

